# Temporal pattern separation in hippocampal neurons through multiplexed neural codes

**DOI:** 10.1101/421479

**Authors:** Antoine D. Madar, Laura A. Ewell, Mathew V. Jones

**Author notes:** **Corresponding authors** Antoine D. Madar (,).

## Abstract

Pattern separation is a central concept in current theories of episodic memory: this computation is thought to support our ability to avoid confusion between similar memories by transforming similar cortical input patterns of neural activity into dissimilar output patterns before their long-term storage in the hippocampus. Because there are many ways one can define patterns of neuronal activity and the similarity between them, pattern separation could in theory be achieved through multiple coding strategies. Using our recently developed assay that evaluates pattern separation in isolated tissue by controlling and recording the input and output spike trains of single hippocampal neurons, we explored neural codes through which pattern separation is performed by systematic testing of different similarity metrics and various time resolutions. We discovered that granule cells, the projection neurons of the dentate gyrus, can exhibit both pattern separation and its opposite computation, pattern convergence, depending on the neural code considered and the statistical structure of the input patterns. Pattern separation is favored when inputs are highly similar, and is achieved through spike time reorganization at short time scales (< 100 ms) as well as through variations in firing rate and burstiness at longer time scales. These multiplexed forms of pattern separation are network phenomena, notably controlled by GABAergic inhibition, that involve many celltypes with input-output transformations that participate in pattern separation to different extent and with complementary neural codes: a rate code for dentate fast-spiking interneurons, a burstiness code for hilar mossy cells and a synchrony code at long time scales for CA3 pyramidal cells. Therefore, the isolated hippocampal circuit itself is capable of performing temporal pattern separation using multiplexed coding strategies that might be essential to optimally disambiguate multimodal mnemonic representations.

**Author Summary:** Pattern separation (the process of disambiguating incoming patterns of neuronal activity) is a central concept in all current theories of episodic memory, as it is hypothesized to support our ability to avoid confusion between similar memories. For the last thirty years, pattern separation has been attributed to the dentate gyrus of the hippocampus, but this has been hard to test experimentally. Moreover, because it is unclear how to define activity patterns in the brain, such a computation could be achieved in many different ways. Here, we demonstrate that pattern separation is performed by hippocampal networks (dentate gyrus and CA3) through a variety of neural codes. By systematically testing different definitions of what it means for spike trains to be similar (using a range of time scales and various standard and innovative metrics that assume different views of the neural code), we assessed how the input-output transformation of multiple hippocampal celltypes relate to pattern separation and found that different celltypes favor complementary coding strategies. This might help storing rich but concise and unambiguous representations of complex events. Finally, we provide the first experimental evidence of the importance of inhibitory signals in mediating pattern separation, and identify through which coding strategies.

## Introduction

Understanding what computations hippocampal neurons perform and how they support episodic memory is a longstanding goal in neuroscience. The hippocampus is notably critical for the discrimination of memories that are similar in content [1]. This cognitive function has long been hypothesized to be supported by a neural process called pattern separation, which is generally thought to be implemented in the dentate gyrus (DG) [2-4]. Pattern separation is defined as the transformation of a non-simultaneous set of similar input patterns of neuronal activity into less similar output patterns [5], and is theorized to happen in the hippocampus before encoding a memory trace in order to avoid confusion between similar memories [6]. Despite a long history of research on the subject, mostly in silico [7], it is still unclear how the activity of single neurons underlies this computation.

Generally speaking, single-neuron computations have been investigated experimentally using two broad strategies: 1) by determining how the spiking of individual neurons, recorded in vivo, is tuned to different parameters of the behavior or environment [8] or 2) by relating the neuronal infra or suprathreshold membrane potential responses, generally recorded in vitro, to their synaptic inputs.

The first strategy has led to the discovery that the principal cells of the hippocampus fire preferentially in specific locations of an environment called place fields [9]. Upon sufficient modification of the environment, the place field(s) of a given neuron remap by relocating or changing their firing rate (in extreme cases, disappearing) [10, 11]. These different forms of remapping have often been used as a proxy for pattern separation, yielding conflicting interpretations on the role of DG and CA3 [12-14]. However, neurons remapping between two similar environments is not sufficient evidence that those neurons participate in pattern separation because place field remapping is assessed without knowledge of the direct synaptic inputs: it is possible that separated representations are inherited from upstream networks. Thus, despite extensive recordings of large populations of several hippocampal celltypes during behavior [11, 14], it remains unknown how each celltype participates in pattern separation.

In contrast to the first strategy, the second one explicitly takes into account the synaptic inputs of recorded neurons. Such an approach has been used to study dendritic integration [15], short [16, 17] or long-term plasticity [18, 19] and the relationship between excitatory inputs and the output spiking probability [19-23], but most of these studies were limited to simple input activity patterns (i.e. rhythmic series of spikes or bursts of spikes) and simple output patterns (i.e. a single spike). Some have used more complex sequences of spatially distributed synaptic inputs [24-26], but the resulting input patterns were still spikes regularly spaced in time. Yet, neuronal spike trains recorded in vivo are not so regular but instead approximately follow a Poisson distribution [27] or a more bursty distribution [28, 29], and what neurons integrate is thus a barrage of irregular synaptic inputs [30, 31]. To infer the single-neuron suprathreshold input-output relationship in more naturalistic regimes, some have thus recorded the spiking response to intracellular injections of current waveforms resulting from either white noise [32] or from the simulated random discharges of a large number of presynaptic neurons [27, 30]. But this only gives insight on the operations performed by a neuron isolated from its network. To understand the role of the network, rare studies have directly stimulated afferents with naturalistic spike trains to characterize short-term [33-35] or long-term [35, 36] synaptic dynamics of hippocampal neurons. An even smaller number of studies have aimed at characterizing the suprathreshold input-output function of a neuron in response to external Poisson spike trains [37, 38], and such experiments have determined that this function was different when evaluated with simpler input patterns [39], confirming the necessity to assess neuronal computations with complex, naturalistic input patterns.

Surprisingly, because none of the above studies have quantified and systematically varied the similarity between input patterns, none have directly addressed how the input-output transformation in single hippocampal neurons relates to pattern separation. To answer this question, we developed a new paradigm [40], in mouse brain slices, where afferent axons of the DG are stimulated with complex input spike trains of varying similarity while the output spike trains from a single neuron are recorded. Thus, in contrast to in vivo studies, we considered input and output temporal patterns instead of spatial ones. With this paradigm, we recently demonstrated that the suprathreshold responses of the principal neurons of the DG and CA3 were strongly decorrelated compared to their inputs, whereas DG interneurons exhibited less temporal decorrelation [40].

Importantly, our previous investigation only considered one type of neural code (binwise synchrony, as measured by the Pearson’s correlation coefficient), even though temporal pattern separation could, in principle, be achieved in a variety of ways depending on the features considered relevant in a spike train. For example, if the only feature carrying information in a spike train was its firing rate, a pattern separator would convert a series of input spike trains with similar rates into a series of output spike trains with dissimilar rates. Alternatively, the firing rate could be irrelevant and two output trains with the same number of spikes could be considered separated if their spiketimes are desynchronized. Pattern separation is thus a group of potential computations, each computation corresponding to a specific neural code, or, in other words, to a specific definition of “similarity”. By testing a wide range of similarity metrics and time scales, the present work aims to determine through which neural codes temporal pattern separation is performed in the hippocampus.

## Results

### Temporal pattern separation via decorrelation, orthogonalization and scaling

In Madar et al. (2018) [40], we demonstrated for the first time that single GCs exhibit high levels of pattern decorrelation thanks to a novel pattern separation assay in acute brain slices. Our general paradigm has three steps (**Fig 1, Methods - Experiments**): 1) ensembles of stimulus patterns (simulating trains of action potentials) are generated, with known degrees of similarity to each other. These sets of input spike trains are then fed into the DG by stimulating the lateral perforant path. 2) The simultaneous response of a single neuron is recorded in whole-cell current-clamp. 3) The similarity between the output spike trains is compared to the similarity between the input patterns, revealing the degree of separation or convergence.

**Fig 1.**
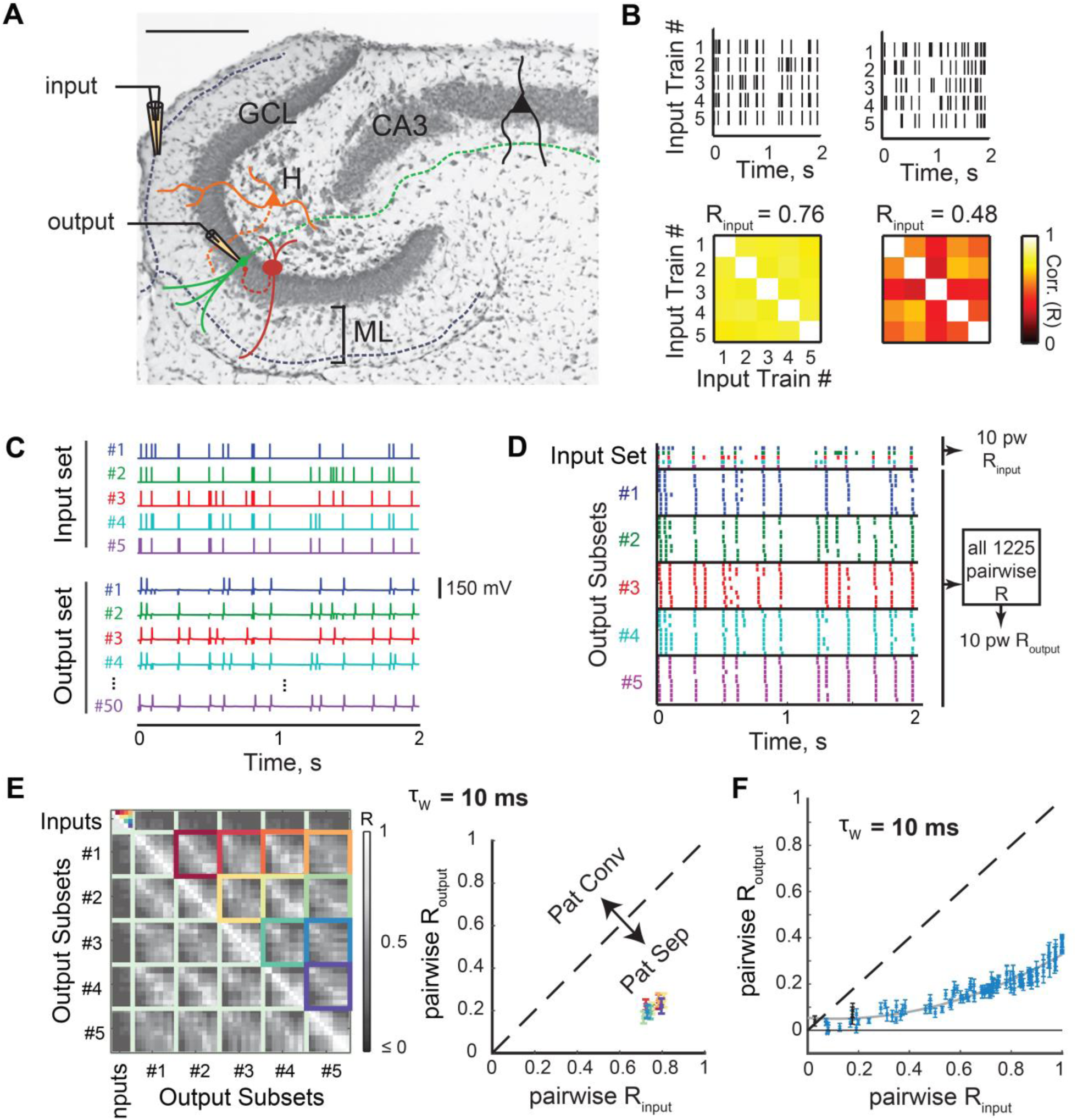
Temporal pattern decorrelation at the level of single dentate granule cells. **(A)** Histology of the DG in a horizontal slice (Cresyl violet/Nissl staining; scale bar: 250 µm), overlaid with a schematic of the experimental setup: a theta pipette in the ML is used to focally stimulate the perforant path (input) while a responding GC is recorded via whole-cell patch-clamp (output). GCL: granule cell layer, H: hilus, ML: molecular layer. Celltype color code: green for granule cell (GC); red for fast spiking interneuron (FS); orange for hilar mossy cell (HMC); black for CA3 pyramidal cell. Solid lines represent dendrites and dashed lines axons **(B)** Examples of input sets. *Top*: each input set is constituted of five different trains of electrical pulses following a Poisson distribution and an average rate of 10 Hz. *Bottom*: correlation coefficient matrix for each input set, each square is the Pearson’s correlation coefficient (R) between two input trains considered as vectors of spike counts with a binning window (τ_w_) of 10 ms. R_input_ is the average of coefficients, diagonal excluded. We used 11 such input sets (R_input_ = 0.11 to 1). **(C)** Current-clamp recordings of the membrane potential of a single GC in response to a single input set (average R_input_ = 0.76). Each input set (five input trains) is repeated ten times, yielding a recording set of fifty output voltage traces. **(D)** Sorted input and output rasters from the recording set in C (each parent input train has ten children output trains forming an output subset, with matching colors). PW means pairwise. **(E)** *Left*: Corresponding 55×55 correlation coefficients matrix using a τ_w_ of 10 ms. Each small grey square represents the correlation coefficient between two spike trains. The average of coefficients from children output trains corresponding to one of the ten pairs of input trains (identified by color-coded squares), was computed to yield ten pairwise R_output_ coefficients. This excludes comparisons between outputs generated from the same parent input. *Right*: mean across multiple cells, following the same color-code as displayed in the matrix on the left. **(F)** Pattern separation graph showing the mean ± SEM pairwise R_output_ across 102 recordings (from 28 GCs), for 11 different input sets (i.e. 110 pairwise R_input_). Values under the identity line (dashed) demonstrate that output spike trains were less similar to each other than inputs, and thus that pattern separation was performed. Blue indicates significant difference from the identity line (one-sample t-test on the difference pairwise R_input_ -pairwise R_output_, p < 0.05). The solid grey line is a parabolic fit to the 102 recording sets. [Fig adapted from Madar et al. (2018) [40] (fig. 1, 2, 5)]

We first reanalyzed a dataset, presented in Madar et al. (2018) [40], of GC recordings performed in response to input sets constituted of 10 Hz Poisson trains (**Fig 1B**). A pairwise similarity analysis (**Fig 1E**), with the Pearson’s correlation coefficient R as a measure of similarity, confirmed our finding that the output spike trains of GCs are significantly less correlated than their inputs at both low and high input correlations (**Fig 1F**). As detailed in Madar et al. (2018) [40], even the repetition of the same input train leads to decorrelated output trains, due to the probabilistic nature of synaptic transmission.

We used R as a similarity measure between spike trains because it is easy to implement and is commonly used to quantify the similarity between neural activity patterns, both in computational [41] and experimental studies [12, 13, 42, 43]. However, the original Hebb-Marr framework theorized pattern separation as the orthogonalization of the input patterns [5, 44]. As a result, the terms “decorrelation” (corresponding to output patterns, viewed as variables, with a lower Pearson’s correlation coefficient than their inputs) and “orthogonalization” (corresponding to output patterns, viewed as vectors, that are closer to a right angle than inputs) are often conflated in the literature, even though they are not mathematically equivalent and have a nonlinear relationship to each other (**Fig 2A-C, S1**. See **Methods – Similarity metrics** and **Supplements - 1**). For instance, pairs of spike trains can be uncorrelated (R = 0) without being orthogonal, or can be orthogonal without being uncorrelated (**Fig 2A-C** and **S1**). To determine whether output spike trains of GCs are truly orthogonalized, we explicitly considered spike trains as vectors of spike-counts and computed the normalized dot product (NDP, i.e. the cosine of the angle between two vectors) between pairs of spike trains to assess their similarity (**Fig 2A, C**. See **Methods – Similarity metrics** and **Supplements - 1**). For every recording set, NDP_output_ was lower than NDP_input_, indicating that the angle between output spike trains was closer to a right angle (i.e., closer to orthogonal) than their inputs (**Fig 2D-E**).

**Fig 2.**
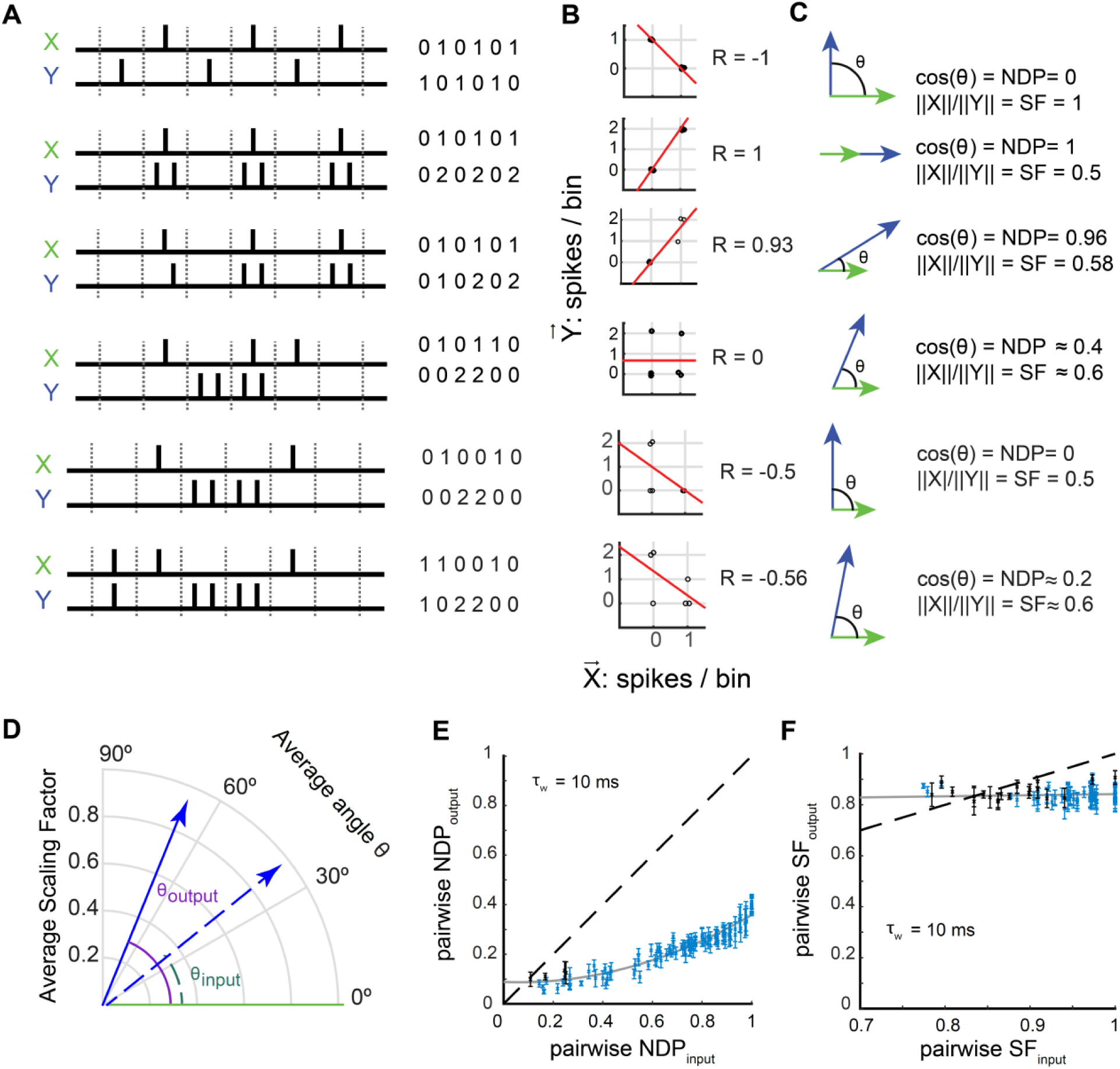
Orthogonalization of input spike trains is a strong component of temporal pattern separation by single granule cells. **(A-C)** Six hypothetical cases of pairs of spike trains and their associated Pearson’s correlation coefficients (R), normalized dot products (NDP, cosine of the angle between two vectors) and scaling factors (SF, ratio of the norms), showing that the three metrics assume different neural codes and are not equivalent. **(A)** Synthetic spike trains (X and Y pairs) divided into six bins, with the corresponding number of spikes per bin. **(B)** R between each pair of X and Y describes the linear regression between the number of spikes in the bins of X versus the corresponding bins in Y (jitter was added to make all points visible). **(C)** Geometric view of vectors X and Y, where each bin is a dimension of a 6-dimensional space, and the number of spikes in a bin is the coordinate along this dimension. NDP measures how far from orthogonal two spike trains are and SF measures how different their binwise firing rates are. (**A-C)** NDP and R are sensitive to whether spike trains have spikes in the same bins or not (row 1), whereas SF is sensitive to differences in spike counts per bin (row 2), although neither NDP nor R is purely about “bin synchrony” (row 3). Note that these examples provide the intuition that orthogonal vectors (NDP = 0) necessarily correspond to a negative correlation between the spike trains (row 1 and 5) but that anticorrelated spike trains (R < 0) are not necessarily orthogonal (row 6), and that orthogonal spike trains are not necessarily perfectly anticorrelated as in row 1 because R, unlike NDP, considers common silent bins as correlated (row 5). See also **Fig S1. (D)** Vector representation of experimental data from one recording set, showing the average similarity between a set of input spike trains (dashed line and green angle, R_input_ = 0.76) and the average similarity between the fifty corresponding output spike trains excluding comparisons between outputs from same parent input (solid line, purple angle). The angles are derived from the NDP whereas the lengths of each vector express differences in binwise firing rates (SF). Here, outputs are more orthogonal (closer to 90°) than their inputs with little difference in scaling. **(E-F)** Pattern separation graphs showing the pairwise output similarity as a function of the pairwise input similarity, as measured by the NDP or SF (mean+/-SEM) across 102 recording sets (28 GCs) for 11 different input sets. As in Fig 1F, blue indicates a significant difference from the identity line (one-sample t-test, p < 0.05). The solid grey line is a parabolic (E) or linear (F) fit to the 102 recording sets. **(E)** Outputs are closer to orthogonality (NDP = 0) than their respective inputs, demonstrating strong levels of pattern separation through orthogonalization at the 10 ms time scale. **(F)** At the 10 ms time scale, pattern separation by scaling is present for inputs very similar in terms of SF (SF_input_ close to 1), but weak, at least over the span of tested SF_input_ values.

Vectors can differ by their angle, but also by their norm. In other words, even if neurons fire in the same time bins (relative to the start of each sweep), the number of spikes per bin can be different, as quantified by the ratio between their norms (scaling factor, SF) (**Fig 2A, C.** See **Methods – Similarity metrics** and **Supplements - 1**). Our results show that for input patterns highly similar in terms of SF, SF_output_ is slightly but significantly lower than SF_input_ (**Fig 2F**). This indicates that variations in the binwise firing rate of single GCs in response to similar inputs is a potential, but weak, mechanism of pattern separation at the 10 ms time scale.

As a whole, these results confirm that input spike trains are decorrelated in the DG at the level of single GCs, and demonstrate that, at the 10 ms time scale, GCs exhibit temporal pattern separation mediated by high levels of orthogonalization and weak levels of scaling.

### Pattern separation of spike trains occurs through multiplexed neural codes

Trains of action potentials can be described in many different ways: their overall firing rate, the timing of their spikes or the number of bursts of spikes, among other statistics (from now on referred to as *spike train features*). Measuring the similarity between two spike trains is not a trivial problem, if only because it is unclear what spike train features are relevant to the brain and what counts as different or similar. In principle, pattern separation could be achieved by the variation of any spike train feature (e.g. if the relevant feature is the total number of spikes in two second bins, pattern separation will be achieved if this number varies more in the set of output spike trains than in the input set). In other words, many different forms of pattern separation could be performed depending on the neural code considered. In our analyses, the neural code (i.e. the set of assumptions on which spike train features are considered relevant and what definition of similarity is used) is determined by the choice of similarity metric and time resolution (see **Supplements – 1**). In order to better characterize which neural code(s) are used to perform temporal pattern separation in the hippocampus, we thus tested multiple metrics and time scales. This approach also allowed us to ask whether several coding strategies could be multiplexed (i.e. simultaneously relevant over different time scales) to achieve pattern separation.

The three measures of similarity we have used above (R, NDP and SF) all carry different assumptions about the neural code (**Table 1)**: R and NDP are mostly, but differently, sensitive to the binwise synchrony whereas SF evaluates variations in spike number (**Fig 2A-C** and see **Supplements - 1**). On the other hand, these metrics resemble each other in that they require binning spike trains in time windows of a prespecified duration (τ_w_). The time scales that are meaningful for the brain are uncertain so we assessed the separation of spike trains for different τ_w_. Indeed, different durations of τ_w_ assume a different window to read the information contained in a spike train. Our analysis shows that pattern separation, measured through R or NDP, is more pronounced at short time scales (e.g. 5 ms) than at longer ones (e.g. 100 ms) (**Fig 3A-B**). In contrast, in the case of high input similarity, pattern separation through scaling is rather weak at short time scales but gets stronger at longer ones (**Fig 3C**). This demonstrates that multiplexed coding allows temporal pattern separation to be carried out by the DG for a large range of time scales.

**Table 1.** Similarity metrics assumption on the neural code

|  | **R** | **NDP** | **SF** | **SPIKE** |
| --- | --- | --- | --- | --- |
| Binned (spike count code) | X | X | X |  |
| Binless (spike time code) |  |  |  | X |
| Preserved time relationship between time points or bins |  |  |  | X |
| "raw" (not baseline-subtracted) spike count vectors |  | X | X |  |
| Assumption of linear relationship between spike trains | X | X | X |  |
| Assumption of Euclidean space | X | X | X |  |
| Excludes common periods of silence |  | X |  |  |
| Mostly sensitive to binwise synchrony | X | X |  |  |
| Sensitive to spike time synchrony |  |  |  | X |
| Sensitive to differences in firing rate or burstiness between spike trains |  |  | X | X |

**Fig 3.**
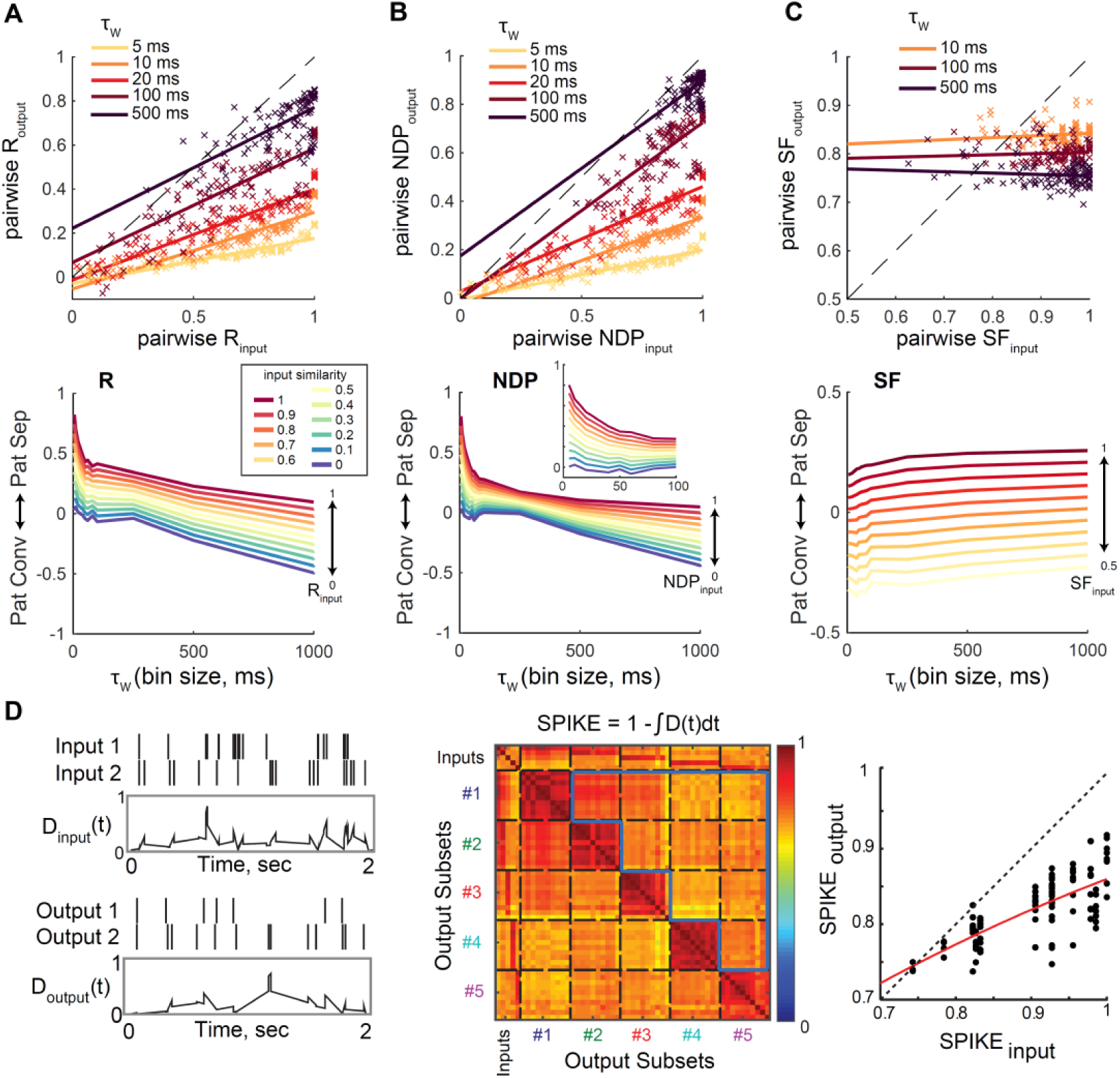
Single granule cells exhibit pattern separation on millisecond to second time scales using different codes. **(A-C)** Pattern separation/convergence assessed at different time scales and using different measures of similarity (S). *Top*: pairwise S_output_ (average across recording sets) as a function of pairwise S_input_, measured with different binning windows τ_w_. Solid curves are linear fits. Each color corresponds to a different τ_w_ ranging from 10 ms to 500 ms. *Bottom*: Effect of τ_w_ on the effective decorrelation (S_input_ – S_output_), interpolated from the linear regressions. Note that as τ_w_ increases, pattern separation through decorrelation (R) or orthogonalization (NDP) becomes weaker while it becomes stronger through scaling (SF). **(D)** Similarity between spike trains is here assessed with the binless SPIKE metric, directly using spike times. *Left*: example of two input spike trains associated with two output spike trains from a GC recording set, and the corresponding distances D(t) between spike trains. D(t) can then be integrated over time to give a single value D. *Middle:* example of 55×55 matrix of SPIKE similarity (1-D) between all spike trains of an example recording set. 0 means that spikes of two trains never happened close in time, and 1 that they were perfectly synchronous. The output SPIKE similarity (SPIKE_output_) is defined as the average similarity excluding comparisons between spike train from the same parent input train (i.e. average of the 10 pairwise SPIKE_output_). *Right*: SPIKE_output_ of the same 102 GC recordings as in Fig 1-3, as a function of SPIKE_input_, fitted with a parabola (red line). Most data points are below the dashed identity line indicating that output spike trains are less similar than inputs. The average SPIKE_input_-SPIKE_output_ is significantly above 0 for all input sets except the two most dissimilar (SPIKE_input_ = 0.74, 0.78) (one-sample T-tests, p < 0.05).

In addition, because a long stream of research suggests that spike trains can carry information directly through the timing of individual spikes [45-47], we also assessed the similarity between spike trains using SPIKE, a binless metric purely based on spike times [48], thus assuming a neural code completely different from the binned metrics above (**Table 1, Methods – Similarity metrics** and **Supplements**). Our results show that input spike trains with similar spike times relative to their sweep start (defined here as spike trains with a high degree of *synchrony*, see **Methods – Similarity metrics**), are transformed into significantly less synchronous outputs, thereby demonstrating that temporal pattern separation in single GCs can occur through spike timing modifications (**Fig 3D**). This is consistent with the high levels of decorrelation and orthogonalization at very short time scales (**Fig 3A-B**) and confirms that temporal pattern separation can be achieved through multiple coding strategies.

### Single GC computations depend on the statistical structure of their input patterns

Results in **Fig 2F** and **3C** show that the similarity of GCs output spike trains in terms of SF is almost flat, suggesting it might be independent of the input similarity, hence leading to significant pattern convergence in certain conditions of low input similarity and short time scales. However, those experiments were based on responses to ∼10 Hz Poisson input spike trains that did not differ much in terms of SF. To further characterize single GC computations, we designed two new input sets spanning a wider range of SF values (**Fig 4A-B**). From its definition, it was apparent that, unlike R and NDP, SF is very sensitive to both differences of overall firing rate and differences in burstiness between spike trains (**Fig S2**). Input set A was thus made of highly correlated Poisson spike trains with different firing rates (7-31.5 Hz), and input set B was made of uncorrelated spike trains with constant firing rate but different levels of burstiness (see **Methods - Experiments**). By recording the spiking output of single GCs in response to these input sets we could thus test how GCs transform input patterns with vastly different statistical structures, and whether they can still perform pattern separation in those conditions. Our results indicate that for both input sets GCs can perform weak levels of pattern separation via scaling for highly similar inputs (**Fig 4C**), consistent with our experiments using 10 Hz Poisson trains (see Fig 2F and 3C). However, although not independent of input similarity, the similarity of GCs output spike trains (in terms of SF) is comprised in a relatively narrow band, yielding high levels of pattern convergence for relatively dissimilar input spike trains (**Fig 4C**). In other words, when assuming a neural code based on the norm of spike count vectors, GCs outputs are maintained relatively similar to each other, even when their corresponding input patterns are very different. This is true from the millisecond to the second time scale, but pattern convergence decreases and separation increases with longer time scales (**Fig 4C**), consistent with previous experiments (see Fig 3C).

**Fig 4.**
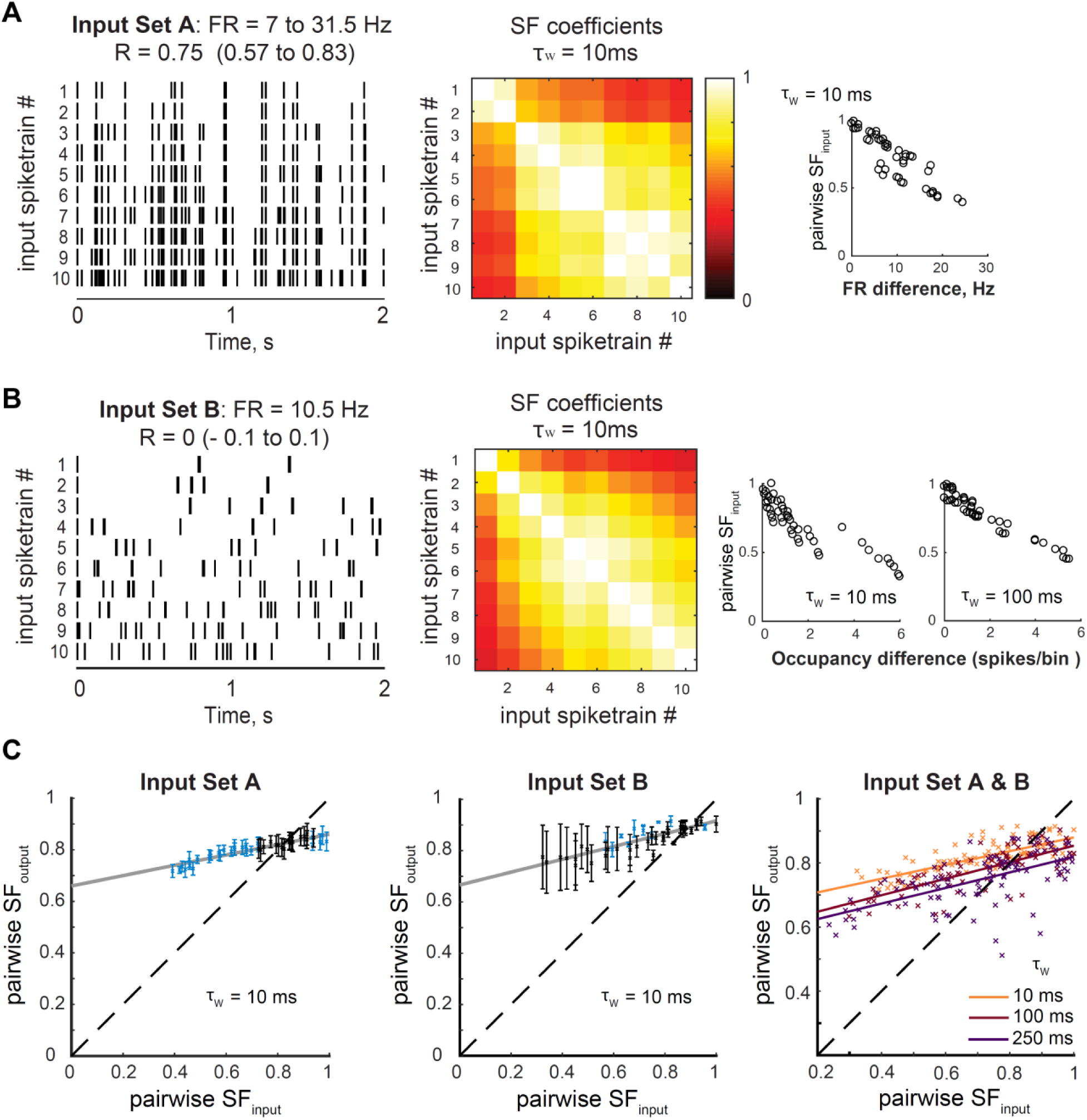
Pattern convergence via scaling in single granule cells. **(A-B)** Two new input sets were designed to explore single GCs computations on inputs with a wide range of pairwise similarity as measured by SF. Both input sets have ten input trains of 2 s, and were repeated five times during the whole-cell patch-clamp recording of a single GC, yielding 50 output spike trains. **(A)** *Left*: Input set A was constituted of spike trains following a Poisson distribution, each with a different firing rate (FR), increased from 7 to 31.5 Hz, but with an average R_input_ constrained around 0.75 (τ_w_ = 10 ms). The ten input trains were thus well correlated but varied in their firing rate. *Middle*: matrix of SF coefficients for each pair of trains in input set A, showing a wide range of values unlike the 11 input sets with 10 Hz trains used in previous experiments. *Right*: SF values are indeed sensitive and anti-correlated to differences in overall firing rate between two spike trains (data points correspond to pairs of trains). **(B)** Because SF is dependent on how differently clustered spikes are in two different trains, SF can vary even when the overall firing rate is the same. *Left*: Input set B was constituted of spike trains with 21 spikes (FR = 10.5 Hz) that were clustered in more or less bins, leading to trains with varying burstiness and thus a wide range of SF values (*Middle*). R was not constrained but ended up close to 0 for all pairs. *Right*: SF values are indeed sensitive and anticorrelated to differences in occupancy (average number of spikes per occupied bins), a direct measure of burstiness when FR is constant across trains (data points correspond to pairs of trains). **(C)** Pattern separation graphs showing the pairwise output similarity as a function of the pairwise input similarity, as measured by SF, across 5 GCs responses to input set A and 3 GCs responses to input set B (Both input sets tested GC responses over 45 pairwise R_input_). *Left and Middle*: Mean+/-SEM. Blue indicates a significant difference from the identity line (one-sample t-test, p < 0.05). The solid grey line is a linear fit to the 5×45 (*Left*) or 3×45 (*Middle*) data points. Both experiments confirm that, at the 10 ms time scale, GCs exhibit pattern separation through scaling for highly similar input trains (SF close to 1) as in Fig 2F, but show that GCs can exhibit pattern convergence for more dissimilar inputs (SF < 0.7). *Right*: Mean across GC recordings for both experiments combined, measured at different time scales. It confirms that SF_output_ decreases when using larger binning windows, leading to more pattern separation for high input similarity and lower pattern convergence for lower input similarities.

Although SF makes mathematical sense when measuring the similarity between vectors, it is difficult to interpret in terms of basic spike train features, in part because it is sensitive to both firing rate and burstiness (see **Supplements – 1** for details). We thus aimed to gain a more intuitive view of how those spike train features vary to result in pattern separation or convergence. For that, we calculated the mean firing rate of each spike train and designed two new indicators providing complementary information on the burstiness of individual spike trains: compactness and occupancy (**Fig 5A**). Briefly, compactness is anticorrelated to the number of time bins occupied by at least one spike (the less occupied bins, the more compact a spike train is), whereas occupancy is the average number of spikes per occupied bin (**Methods – Firing rate and burstiness codes** and **Supplements - 2**).

**Fig 5.**
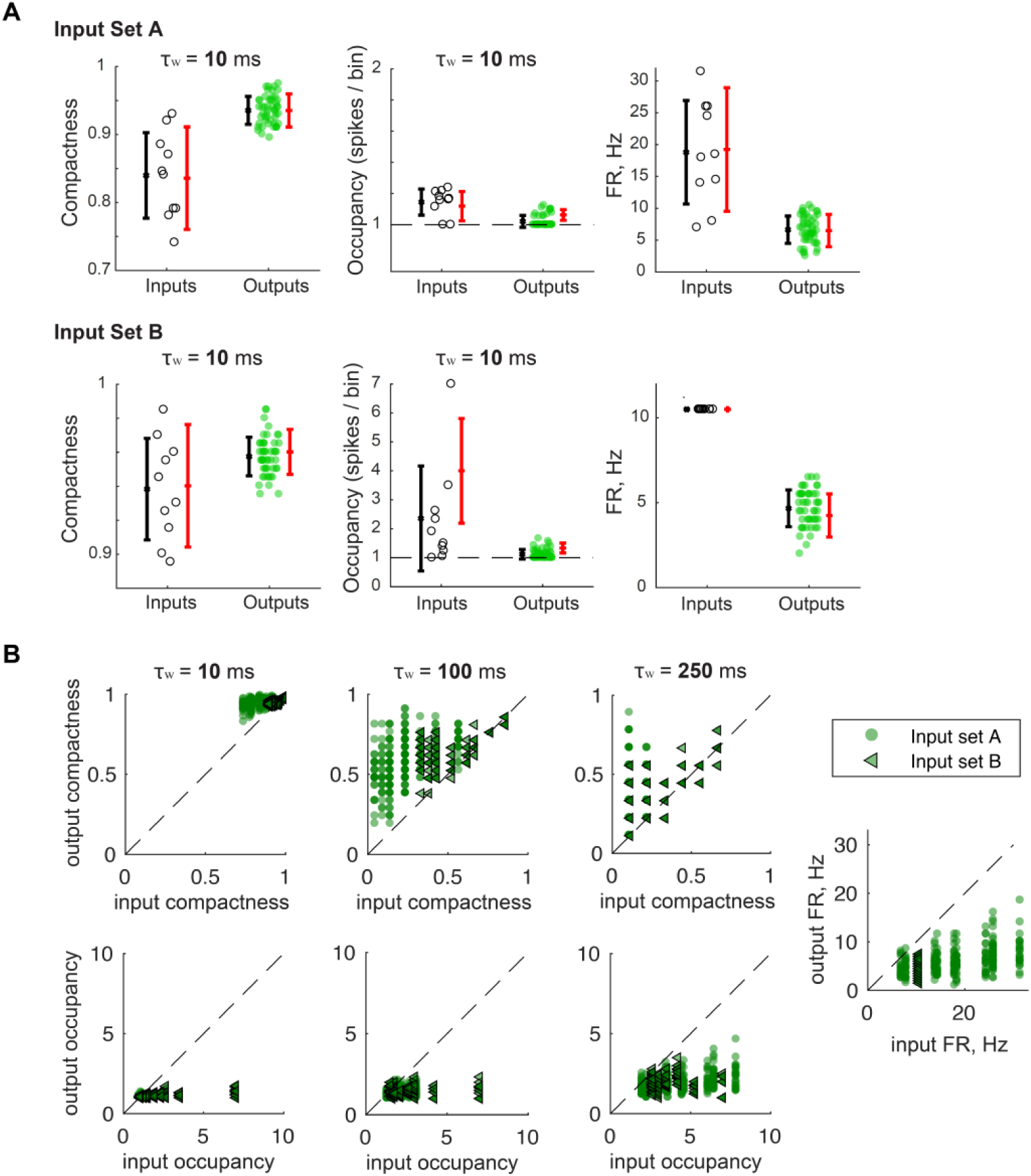
Granule cells input-output transformations in terms of firing rate and burstiness. **(A)** Graphs shown correspond to the same GC in response to either input set A or B. Data points correspond to individual spike trains, (black empty circles for inputs: 10 trains per set; green filled circles for GC outputs: 50 trains per set). For each spike train, we measured its overall FR, binwise compactness and binwise occupancy, all spike train features providing information on sparseness and burstiness (see **Methods**). Error bars correspond to two different measures of dispersion. Black: mean +/- SD; red: min-max center of gravity ± the mean absolute difference (self-comparisons and comparisons between output trains with the same parent input train were excluded). Comparing mean values of a given spike train feature between inputs and outputs shows the direction of the transformation from inputs to outputs. Comparing the dispersion between inputs and outputs shows whether pattern separation (larger output dispersion) or convergence (smaller output dispersion) was achieved through a neural code focused on the measured spike train feature. **(B)** For each output spike train, the output spike train measures of compactness, occupancy or overall FR is plotted against the measure of its parent input train (input set A or B). For all time scales, GCs have generally higher compactness than their inputs, but maintain a low and narrow range of occupancy regardless of their input statistics, showing that GCs are sparser (see **Table S1**). Consistently, GCs maintain a lower FR than their inputs.

First, we asked how GCs transform their inputs in terms of firing rate and burstiness. For all time scales, GCs have generally equal or higher compactness than their inputs but maintain a constant low range of occupancy (**Fig 5B left**). This combination suggests that, using **Table S1**, GC responses are mostly a sparser version of their inputs, with a decrease in the number of spikes per burst when inputs are bursty. Accordingly, the overall firing rate of GC output spike trains is generally lower than their inputs (**Fig 5B right**), which is consistent with the known sparsity of GC activity in vivo and fits the view of the DG acting as a filter of cortical activity [49].

We next asked whether temporal pattern separation is performed through variation of the firing rate between GCs output spike trains, or through variations of compactness or occupancy, or a combination of those three. Our results indicate that pattern separation, but also pattern convergence, can be exhibited for all three types of codes, depending on certain conditions (**Fig 6**). First, separation is favored at longer time scales (for burstiness codes) and high input similarity (for burstiness and rate codes) (**Fig 6A**). Second, the direction of the computation (separation or convergence) strongly depends on the statistical structure of the input patterns (**Fig 6B**): 1) In response to Poisson inputs with very similar firing rates (P10Hz), GCs exhibit low but significant levels of pattern separation through small variations of compactness, occupancy and firing rate, which become more relevant with larger time scales. Pattern separation measured through scaling (see Fig 2F and 3C) was thus due to the variability in all three spike train features combined. 2) In response to Poisson inputs with varying firing rates (PΔFR, input set A) GCs switch, as the time scale is increased, from pattern convergence to pattern separation through compactness variations, while increasing their levels of convergence in terms of occupancy. Accordingly, their output spike trains have more similar firing rates than their inputs (this is because GCs maintain a narrow range of firing rates regardless of the rate of their inputs, as shown in Fig 5B). 3) In response to inputs with a constant firing rate but varying levels of burstiness (B 10.5 Hz, input set B), GCs show low but significant pattern convergence in terms of compactness and occupancy. Pattern convergence in terms of occupancy progressively decreases as the time scale increases. This translates into significant pattern separation through variations of the firing rate (note that, because spike trains were 2 s long, pattern separation via FR variations is equivalent to pattern separation via occupancy variations in a single 2 s time scale). Interestingly, this analysis demonstrates that GCs can exhibit both separation and convergence simultaneously through different neural codes: for instance, at the 250 ms time scale, the compactness of GCs output spike trains vary more than the compactness in input set A, whereas GCs output spike trains occupancy vary less (**Fig 6B, Table 2**).

**Table 2.**
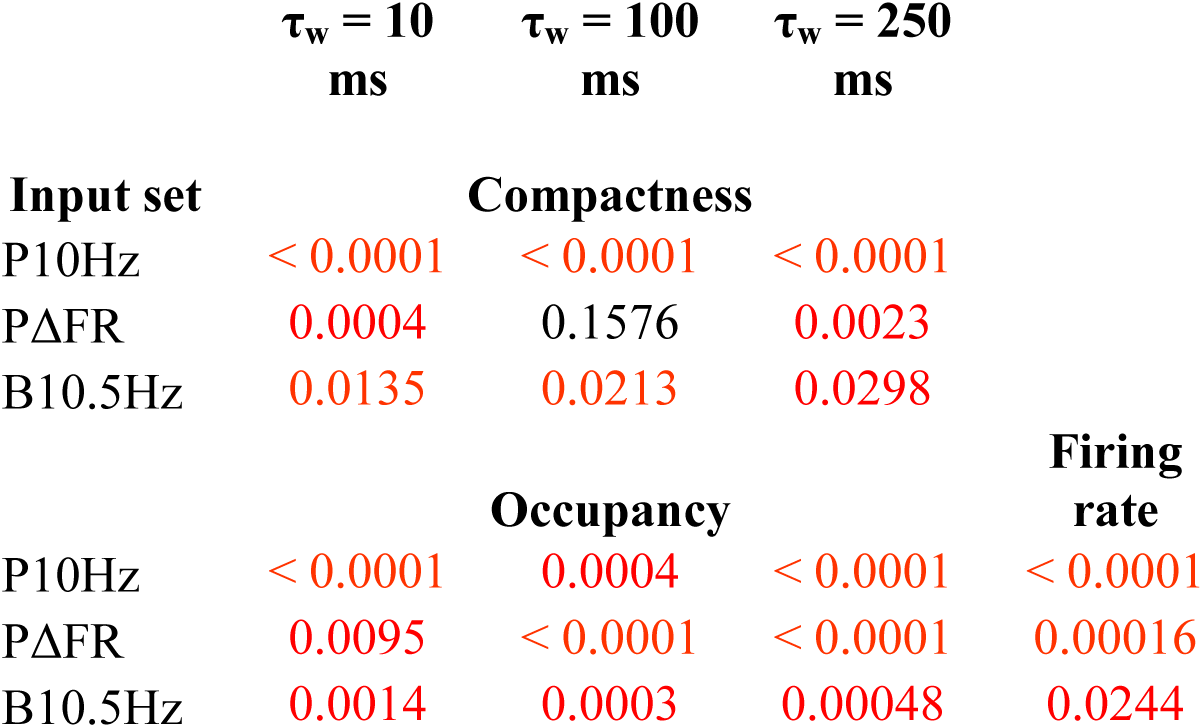
related to Fig 6B One sample t-test (difference from 0) p value < 0.05 in red.

**Fig 6.**
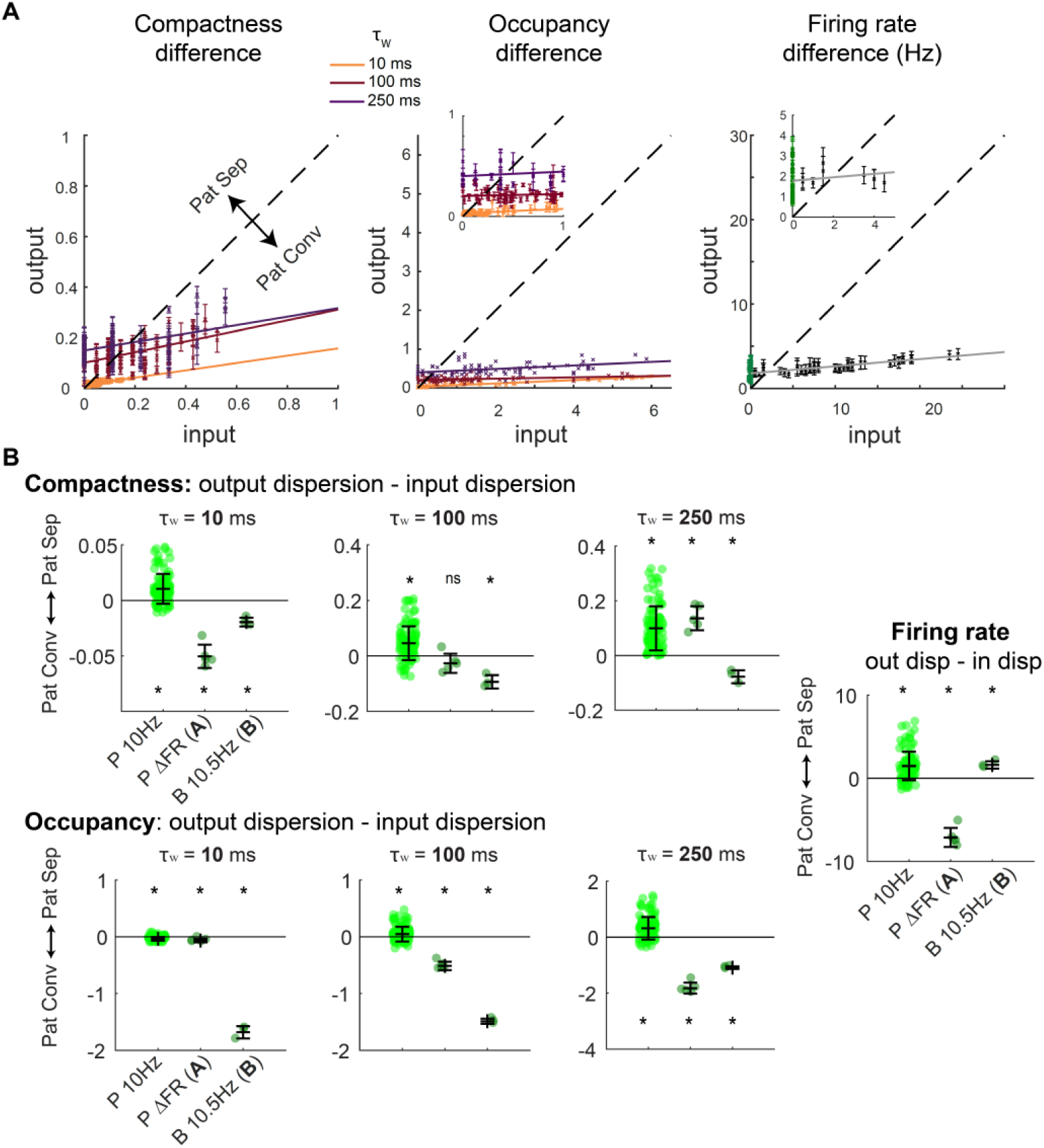
Depending on inputs statistical structure, granule cells exhibit pattern separation or convergence via burstiness and firing rate codes. **(A)** Pattern separation graphs showing the average pairwise output absolute difference across GC recordings as a function of the pairwise input absolute difference, for the binwise compactness (*Left*), binwise occupancy (*Middle*) and overall FR (*Right*). Insets zoom in on the smallest values of the input-axis. Experiments using input set A and B were combined. Note that because we plot the absolute difference between two spike trains (i.e. the distance between two data points in Fig 5A) and not a measure of similarity, pattern separation corresponds to points above the identity line (dashed) and pattern convergence to points below. Error bars are SEM (not displayed in middle graph for readability). *Right*: input set A recordings (PΔFR) in black, input set B (B 10.5 Hz) in green. **(B)** The difference in dispersion (Dispersion_output_ – Dispersion_input_; dispersion is the mean absolute difference, i.e. red bars in Fig 5A) of binwise compactness, binwise occupancy or overall FR for each GC recording set from three experiments using different types of input sets: Poisson trains with 10 Hz FR and varying R (P10Hz: 102 recordings, 28 GCs), Poisson trains with R ≈ 0.75 and varying FR (PΔFR, i.e. input set A: 5 GCs), trains with 10 Hz FR and varying burstiness (B 10.5 Hz, i.e. input set B: 3 GCs). Positive values correspond to pattern separation through variations of the given spike train property (compactness, occupancy or FR) and negative values to pattern convergence. Asterisks denote significant difference from 0, i.e. either pattern separation or convergence (one-sample t-test, p < 0.05. See **Table 2**).

### Inhibition enforces temporal pattern separation in GCs

GCs are known to receive strong feedforward and feedback inhibition from a variety of GABAergic interneurons [50, 51] that impact GCs spiking output [21, 52, 53]. Furthermore, theoretical work has long suggested a role for interneuron interactions with GCs in mediating pattern separation [4, 54, 55], but experimental evidence is lacking. To test the influence of fast inhibitory transmission on the complex computations of GCs characterized above, we recorded GCs responses to 10 Hz Poisson trains before and after application of a nonsaturating concentration of Gabazine, a GABA_A_ receptor antagonist (**Fig 7A**, see **Methods - Experiments**). Under these conditions, IPSC amplitude was reduced by ∼30% (not shown), which led to a slight but visible increase in firing rate, in part due to a higher propensity to fire short bursts of action potentials riding a single EPSP (**Fig 7A-B**). This pharmacological manipulation led to lower levels of temporal pattern separation as measured by R, NDP, but also SF (**Fig 7C**). This was true at all time scales (**Fig 7D**), with the importance of inhibition for pattern separation even increasing at longer time scales. Interestingly, the impact on pattern separation at larger time scale was not only for SF, whose relevance increases with larger τ_w_ (see Fig 3C) but also for R and NDP, whose relevance normally decreases with larger τ_w_ (see Fig 3A-B).

**Fig 7.**
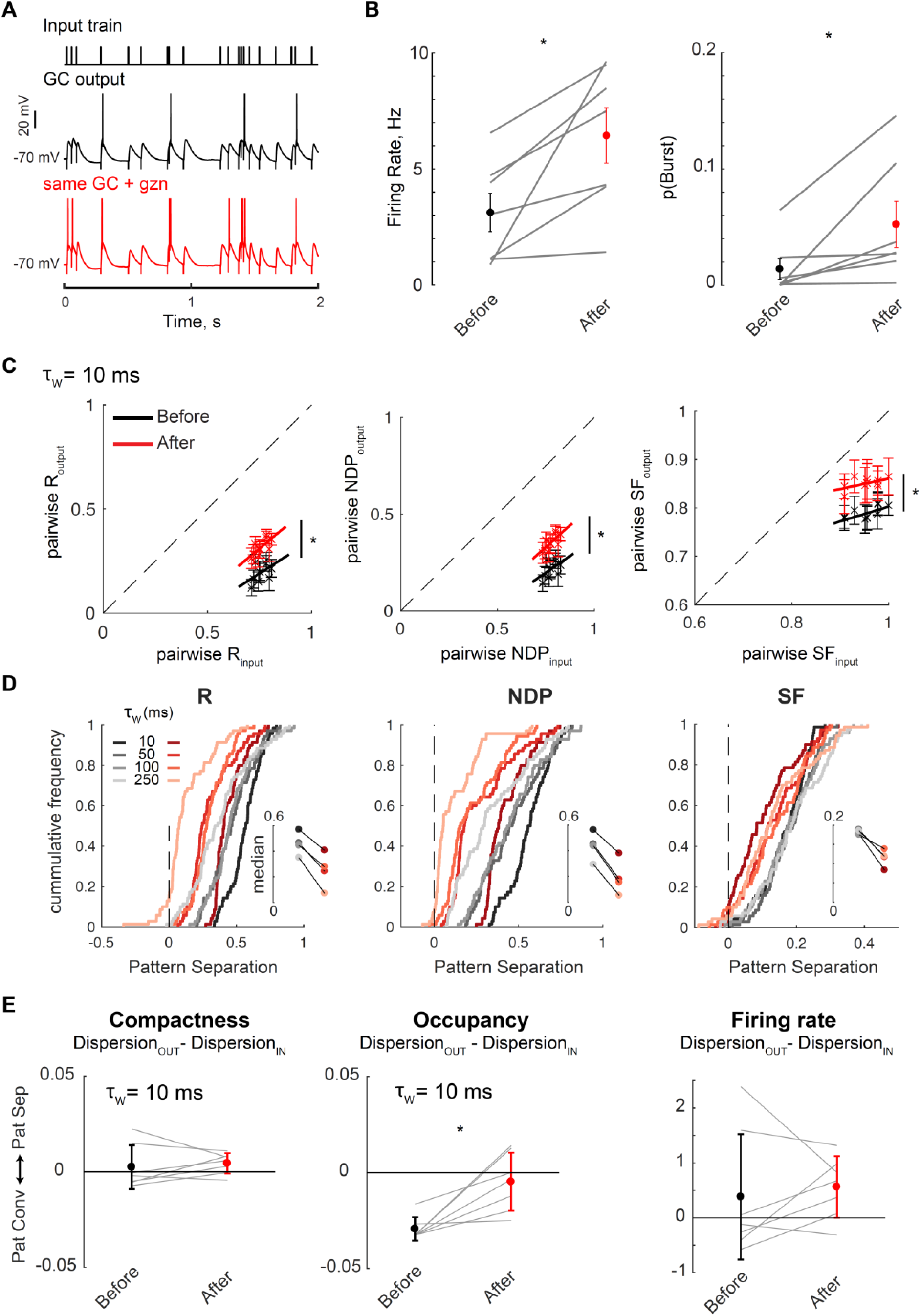
Inhibition controls levels of pattern separation in granule cells through different neural codes. **(A)** Single GCs were recorded twice in response to the same input set (P10Hz, R = 0.76 at τ_w_ = 10 ms, shown in Fig 1B): First in normal aCSF, then under conditions of partial inhibitory block (aCSF + 100 nM gabazine, gzn), without changing any other parameter (i.e. same stimulation location and intensity). **(B)** Partial disinhibition of GCs led to higher FR and a higher propensity to fire small bursts of spikes riding on the same EPSP, as illustrated in A and evidenced by an increased p(Burst) (probability of firing more than one spike between two input pulses). FR and p(Burst) were computed for each recording set (n = 7 GCs). Paired t-test (FR and p(Burst) respectively): p = 0.02, p = 0.04. **(C)** Pattern separation graphs showing the pairwise output similarity as a function of the pairwise input similarity, as measured by R, NDP or SF at the 10 ms time scale (mean+/-SEM) across recordings in 7 GCs. Data points below the identity line (dashed) correspond to pattern separation. Partial block of inhibition in GCs led to a significant decrease in pattern separation through all similarity metrics (ANCOVA with separate-lines model fitting 70 data points per treatment group: p < 0.0001). Solid lines are the linear fits used for the ANCOVA. **(D)** Cumulative frequency distributions of the distance of (pairwise S_input_, pairwise S_output_) data points to the identity line in pattern separation graphs as in C (insets show medians). Positive values of the x-axis correspond to pattern separation, and negative values to pattern convergence. Red curves (after gzn) are all shifted to the left of their black counterparts (before gzn), showing a decrease in pattern separation at all time scales. The shift is more pronounced at larger time scales for R and NDP, at smaller time scales for SF. ANCOVA on (pw S_input_, pw S_output_) data points with separate-lines model: p < 0.0001 for τ_w_ up to 1000 ms for R and NDP, p < 0.02 for SF. **(E)** Average levels of pattern separation via compactness, occupancy or FR codes were measured for each recording set (as in Fig 6B). Surprisingly, partial block of inhibition did not significantly change the variations in FR or binwise compactness of output spike trains, but it led to less pattern convergence or even pattern separation through variations of occupancy (paired t-test: p = 0.008, 0.024, 0.031 for τ_w_ = 10, 20 and 50 ms respectively. No significant difference between treatments for larger τ_w_ or other spike train features at any τ_w_.

The average R_output_ was well correlated to the average firing rate of a recording set (linear regression: R^2^ = 94%, p < 0.0001, n = 14 data points from 7 GCs), suggesting that the inhibitory control of sparsity in GCs firing helps temporal pattern separation as measured with R. Interestingly, in a previous report we did not detect such a strong relationship between R and the firing rate levels of GCs [40]. Moreover, R is mathematically independent of proportional increases of pairwise firing rate levels and not very sensitive to non-proportional ones (see **Supplements - 1** and **Fig S1C**). In the present experiment, the correlation between R and the firing rate is thus likely to arise from physiological reasons that conjointly affect both the sparsity and the binwise synchrony of output spike trains rather than from the mathematical assumptions behind R.

In contrast to R and NDP, the mathematical definition of SF makes it sensitive to pairwise firing rate levels (i.e. SF systematically increases for pairs of spike trains with higher firing rates. See **Supplements - 1** and **Fig S1C**). As seen before, SF is also designed to be sensitive to differences in firing rate and burstiness of individual spike trains (e.g. Fig 5 and S2). Therefore, the effect of gabazine on decreasing pattern separation as measured by SF (**Fig 7C-D**) could be due: 1) to a global increase in firing rates in all the output spike trains, 2) to variations of firing rate or burstiness between output spike trains, or 3) to a combination of all of the above. The direct assessment of pattern separation in terms of firing rate or burstiness indicates that inhibition actually does not affect temporal pattern separation through rate or compactness codes and favors pattern convergence through occupancy codes (**Fig 7E**). We can thus conclude that the effect of inhibition on pattern separation via scaling (SF) is mostly due to the global firing rate increase under gabazine.

Overall, we provide here the first experimental evidence that fast inhibitory transmission in the DG controls levels of temporal pattern separation and convergence through different multiplexed neural codes.

### Different hippocampal celltypes support pattern separation through different neural codes

GCs are embedded in a network of multiple celltypes all participating in generating the output of the DG. For instance, computational models of the DG have suggested that different populations of inhibitory and excitatory interneurons could modulate certain forms of pattern separation [43, 54]. To develop a comprehensive understanding of the computations performed by the DG, and hippocampal networks in general, it is important to characterize the input-output transformation of a diverse set of celltypes. We recorded from dentate fast-spiking GABAergic interneurons (FSs), glutamatergic hilar mossy cells (HMCs) and CA3 pyramidal cells (PCs) in response to 10 Hz (for GC, FS, HMC) or 30 Hz (for PC and GC) Poisson input trains with varying levels of correlation (**Fig 8A** and **Methods – Experiments**).

**Fig 8.**
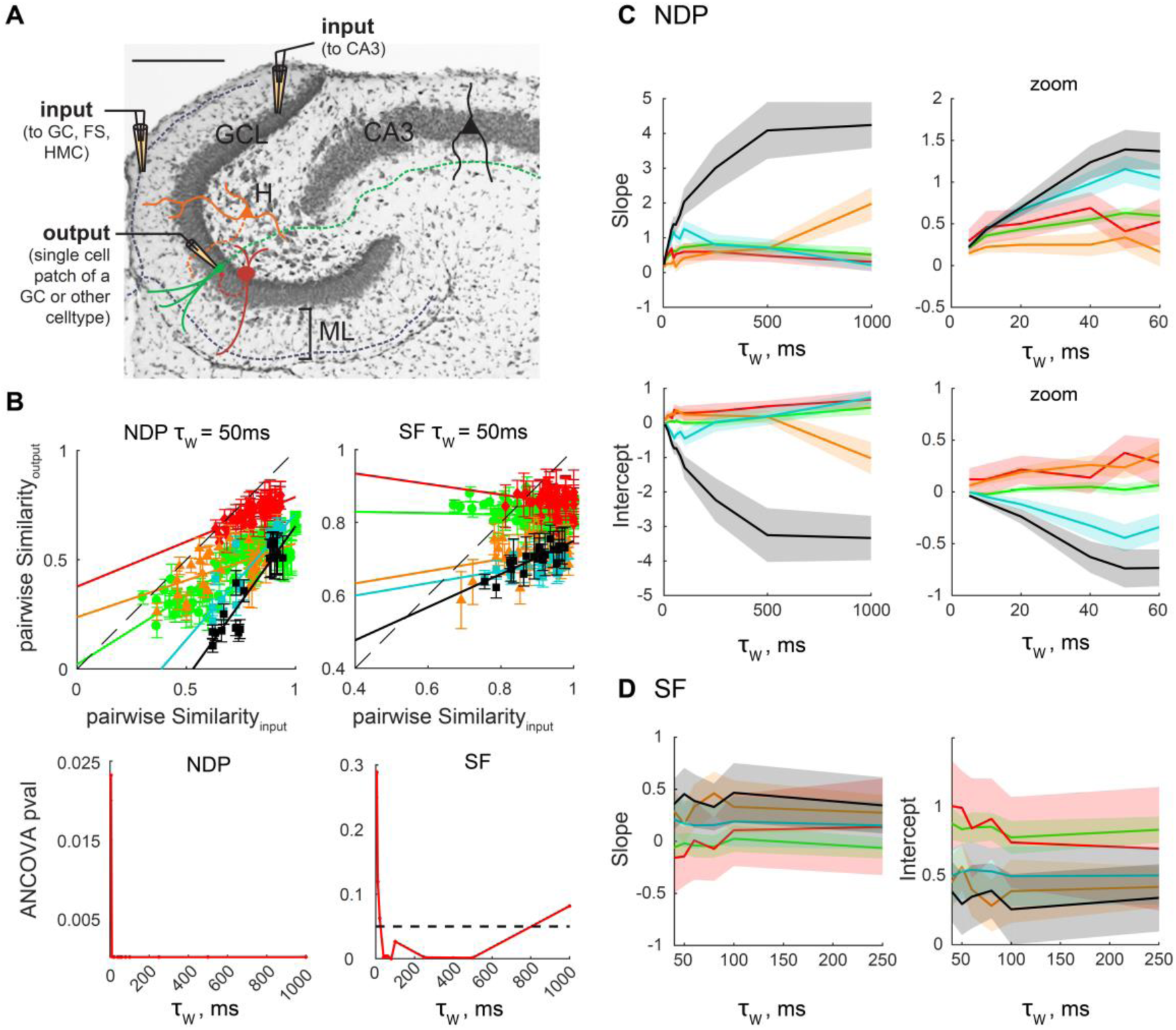
Levels of pattern separation through orthogonalization and scaling differ between hippocampal celltypes. **(A)** Temporal pattern separation assays were performed by single whole-cell current-clamp recordings from GCs (green, same experiments as Fig 1-3), FSs (red), HMCs (orange) and CA3 PCs (black). For GCs, FSs and HMCs, input sets with Poisson trains of 10 Hz FR (e.g. Fig 1B) were delivered by stimulating in the outer molecular layer (OML), and recordings done in regular aCSF. For CA3 PCs those stimulus parameters rarely if ever elicited spiking, therefore input sets were constituted of Poisson trains with a 30 Hz FR delivered in the GCL, and recordings were performed under partial block of inhibition (100 nM Gabazine). Control experiments (teal) were performed in GCs under the same pharmacological conditions using 30 Hz input sets, but the stimulations were delivered in the OML. The membrane potential was maintained around −70 mV (between −70 and −60 mV for CA3 PCs). Note: although all celltypes and treatments are displayed on the same graphs for concision (B-D and Fig 9), CA3 PCs should only be directly compared to their corresponding GC controls. **(B)** *Top*: Pattern separation graphs showing the pairwise output similarity as a function of the pairwise input similarity, as measured by NDP or SF at the 50 ms time scale (mean+/-SEM: 28 GCs, 3-13 recordings per input set; 4 FSs, 4 recordings per input set; 11 HMCs, 5-7 recordings per input set; 14 CA3 PC + gzn, 6-9 recordings per input set; 13 control GCs + gzn, with 11 recordings per input set). Same color code as in A. Data points below the identity line (dashed) correspond to pattern separation. All celltypes performed pattern separation through both orthogonalization and scaling but with different levels. *Bottom*: We performed ANCOVAs with separate-lines model for all five celltypes (e.g. solid lines shown in the top graphs) for 11 different time scales (5-1000 ms). Dashed horizontal line represents significance level at 0.05: orthogonalization and scaling levels differ between celltypes at most time scales. **(C-D)** Slope and intercept of the linear models (e.g. solid lines in B, top) used to fit [S_input_, S_output_] data points for a given celltype and time scale. Shaded patches correspond to the 95% confidence interval of the parameter estimate. Significant differences between celltypes in pattern separation functions (with Tukey-Kramer correction for multiple comparisons) are detailed in **Table 3**.

Previously, we found that temporal pattern separation, measured using R, was not specific to GCs but that GCs exhibit the highest levels of decorrelation among tested dentate neurons, and that CA3 PCs express even more temporal decorrelation than GCs [40]. Here we ask whether this is still the case when considering other neural codes based on different similarity metrics. A pairwise similarity analysis and a comparison of the pattern separation function (the linear regression model describing S_output_ as a function of S_input_, with S representing a given similarity metric) across celltypes shows that pattern orthogonalization (NDP) and scaling (SF) levels significantly differ at most time scales (**Fig 8B-D, Table 3**).

**Table 3.** related to Fig 8C-D Post-hoc t-tests p-values with correction for multiple comparisons when necessary (Tukey-Kramer) comparing slope or intercept estimates (from ANCOVA) between celltypes. Note that CA3+gzn and GC+gzn are only compared with each other. P-values < 0.05 in red.

| Compared Celltypes | | $\tau_w = 10$ ms | $\tau_w = 50$ ms | $\tau_w = 100$ ms | $\tau_w = 250$ ms | $\tau_w = 1000$ ms |
| --- | --- | --- | --- | --- | --- | --- |
|  |  | NDP |  |  |  |  |
| GC | FS | slope: 0.3197,<br>intercept: 0.0205 | slope: 0.2177,<br>intercept: 0.0028 | slope: 0.7884,<br>intercept: 0.2455 | slope: 0.6601,<br>intercept: 0.3798 | slope: 0.8479,<br>intercept: 0.8847 |
| GC | HMC | slope: 0.0038,<br>intercept: <0.0001 | slope: 0.0013,<br>intercept: 0.0009 | slope: 0.0140,<br>intercept: 0.0177 | slope: 0.0758,<br>intercept: 0.0818 | slope: <0.0001,<br>intercept: <0.0001 |
| FS | HMC | slope: 0.0069,<br>intercept: 0.9758 | slope: 0.8737,<br>intercept: 0.4558 | slope: 0.6957,<br>intercept: 0.9756 | slope: 0.9990,<br>intercept: 0.8882 | slope: 0.0001,<br>intercept: <0.0001 |
| GC+gzn | CA3+gzn | slope: 0.8228,<br>intercept: 0.0110 | slope: 0.0920,<br>intercept: 0.0084 | slope: 0.0004 ,<br>intercept: <0.0001 | slope: <0.0001,<br>intercept: <0.0001 | slope: <0.0001,<br>intercept: <0.0001 |
| Compared Celltypes |  | SF |  |  |  |  |
| GC | FS | slope: 0.7487,<br>intercept: 0.6242 | slope: 0.7583,<br>intercept: 0.6209 | slope: 0.9279,<br>intercept: 0.9838 | slope: 0.7575,<br>intercept: 0.9719 | slope: 0.5867 ,<br>intercept: 0.5271 |
| GC | HMC | slope: 0.4701,<br>intercept: 0.1568 | slope: 0.1848,<br>intercept: 0.0138 | slope: 0.0749,<br>intercept: 0.0069 | slope: 0.0028,<br>intercept: <0.0001 | slope: 0.970,<br>intercept: 0.0052 |
| FS | HMC | slope: 0.3924,<br>intercept: 0.1585 | slope: 0.2396,<br>intercept: 0.0477 | slope: 0.6326,<br>intercept: 0.2795 | slope: 0.8893,<br>intercept: 0.5867 | slope: 0.3993,<br>intercept: 0.2901 |
| GC+gzn | CA3+gzn | slope: 0.1092,<br>intercept: 0.1247 | slope: 0.0969,<br>intercept: 0.1231 | slope: 0.1436,<br>intercept: 0.1573 | slope: 0.2810,<br>intercept: 0.3182 | slope: 0.1605,<br>intercept: 0.1942 |

For NDP, we notice a progression from FSs to HMCs to GCs to CA3 PCs, with GCs performing the highest levels of separation among DG neurons, and CA3 PCs surpassing GCs, especially with their ability to orthogonalize inputs already relatively dissimilar (**Fig 8B**). The orthogonalization function of GCs, HMCs and CA3 PCs diverge as the time scale is increased (**Fig 8C**).

For SF, scaling functions do not depend much on the input similarity, except for CA3 PCs. Mostly based on the scaling functions intercept, we notice two groups (**Fig 8D**): FSs and GCs exhibit low levels of pattern separation through scaling, in contrast to HMCs and CA3 PCs (and their GC controls) that produce spike trains with lower similarities in terms of SF. Note that among celltypes tested with 10 Hz Poisson trains, HMCs are the best at pattern separation through scaling, not GCs.

In contrast to GCs, HMCs and FSs tend to fire bursts of spikes (although the characteristics of bursts differ between the two celltypes), which affects, at least partially, their levels of pattern decorrelation [40]. An analysis of the distribution of inter-spike-intervals in all recorded celltypes confirmed that HMCs and FSs are bursty (as they diverge from a Poisson process) but not to the same extent (**Fig 9A,** and see **Methods - Firing rate and burstiness codes**). It also revealed that, although neither CA3 PCs nor their GC controls (recorded under gabazine and 30 Hz inputs) displayed bursts elicited by a single input pulse [40], they can be considered bursty in the sense that their output spikes tend to be clustered together in time in a nonrandom fashion (**Fig 9A**). We thus asked whether these celltypes with diverse bursty behaviors could perform pattern separation through compactness, occupancy or firing rate codes. Our analysis shows indeed that these features vary across the output spike trains of hippocampal neurons, but in different ways for each celltype (**Fig 9B, Table 4**): for example, at long time scales FSs and HMCs achieve pattern separation mostly through variations of occupancy and firing rate, whereas CA3 PCs achieve pattern separation mostly through variations of compactness.

**Table 4.** related to Fig 9B Posthoc t-test after Kruskal-Wallis one-way ANOVA, with Tukey-Kramer correction for multiple comparisons. CA3+gzn and GC+gzn are only compared with each other. P values < 0.05 in red.

| | | $\tau_w = 10 \text{ ms}$ | $\tau_w = 100 \text{ ms}$ | $\tau_w = 250 \text{ ms}$ | | |
| --- | --- | --- | --- | --- | --- | --- |
| Compared Celltypes |  | Compactness |  |  |  |  |
| GC | FS | <0.0001 | 0.4368 | 0.0339 |  |  |
| GC | HMC | 0.0064 | 0.9214 | 0.7015 |  |  |
| FS | HMC | 0.6255 | 0.437 | 0.0326 |  |  |
| GC+gzn | CA3+gzn | 0.0728 | 0.0112 | 0.3864 |  |  |
|  |  | Occupancy |  |  | FR |  |
| GC | FS | < 0.0001 | < 0.0001 | < 0.0001 | <0.0001 |  |
| GC | HMC | 0.0044 | < 0.0001 | 0.1911 | 0.067 |  |
| FS | HMC | 0.6787 | 0.7675 | 0.0467 | 0.1138 |  |
| GC+gzn | CA3+gzn | 0.1732 | 0.8045 | 0.4212 | 0.0592 |  |

**Fig 9.**
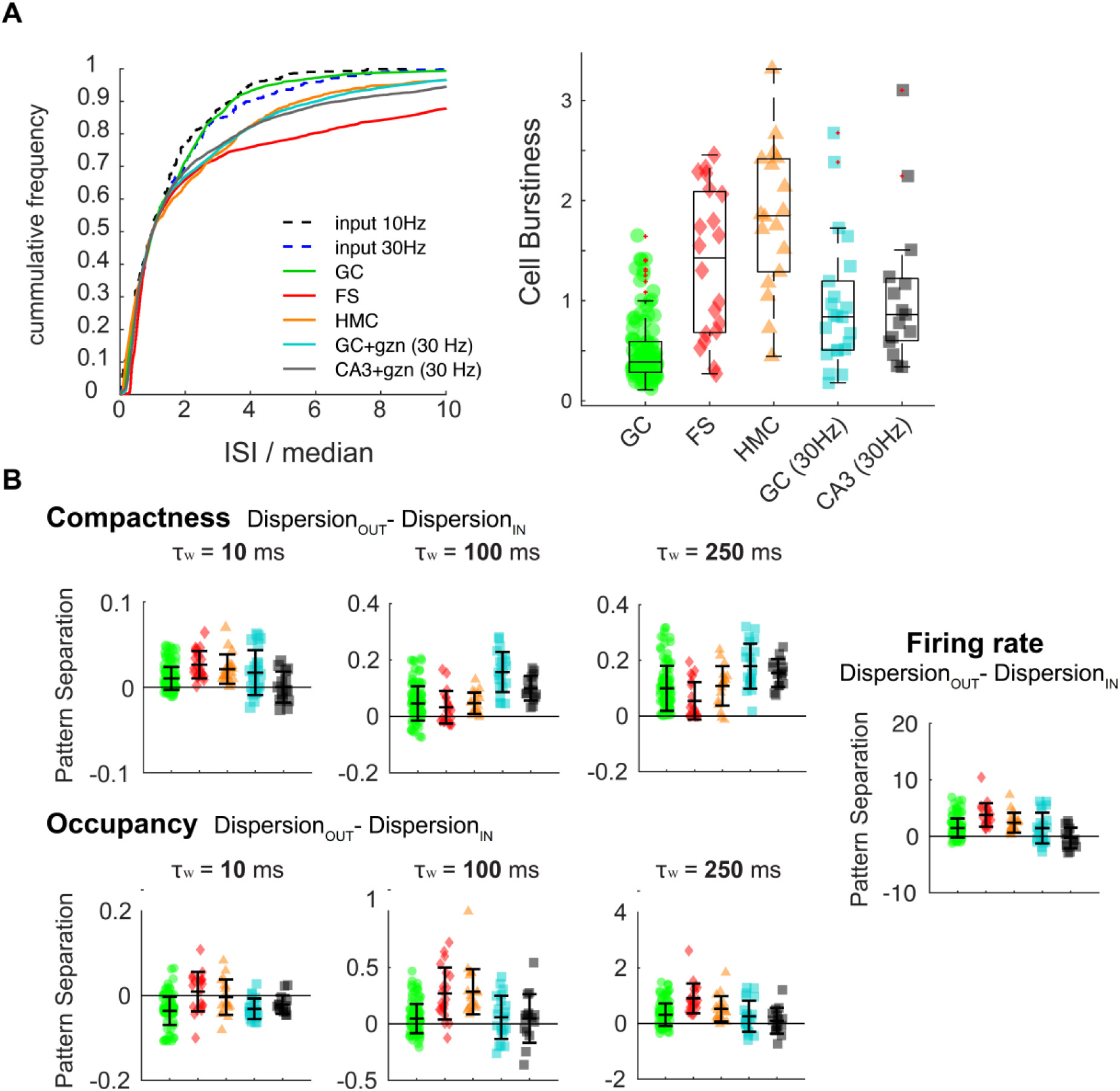
Levels of pattern separation and convergence through burstiness and firing rate codes differ between hippocampal celltypes. **(A)** Assessment of burstiness in different celltypes (binless analysis). *Left*: Cumulative frequency distributions of interspike intervals (ISI) observed in the output spike trains of different hippocampal celltypes and their associated input trains. ISI of all recordings of a given celltype or input set type were pooled. The ISIs were normalized to their median for a given celltype or input type, such that their cumulative distribution would cross at frequency = 0.5, and that all exponential cumulative distributions characteristic of a Poisson process (as both type of input sets are) would be super-imposed or very close to each other. Thus, GCs are visually close to an exponential distribution, suggesting their output is close to a Poisson process, i.e. not bursty. Distributions falling away from their corresponding input distribution suggest that spikes are organized in a non-Poisson manner, suggestive of a bursty behavior. In this sense, HMCs, CA3 PCs and their GC controls seem to exhibit a similar bursty behavior, and FSs an even more bursty one. *Right*: Quantification of the burstiness of each recording set (102 for GCs, 20 for FSs, 18 for HMCs, 15 for CA3 PCs+gzn and 22 for GCs+gzn). Burstiness was assessed as the Kullback-Leibler divergence from P to M, with M the frequency distribution of normalized ISI in an output set (50 output trains), and P the frequency distribution of normalized ISI in the associated input set type (either Poisson 10Hz or Poisson 30 Hz input sets were combined, as in A). The higher the divergence, the burstier a cell is. This analysis confirms that GCs outputs are close to a Poisson process, and that all other celltypes and conditions are significantly burstier (Kruskal-Wallis one-way ANOVA: p < 0.0001; post-hoc t-tests with Tukey-Kramer correction for multiple comparisons: p < 0.01 for all GC (10Hz) to other celltypes comparisons. CA3 PCs and GC controls (30Hz) are not significantly different (p = 0.99) nor FSs and HMCs (p = 0.64)). Box plots: central mark and edges are the median, the 25th and the 75th percentiles, respectively. Whiskers are most extreme data points not considered outliers (red +). **(B)** Average levels of pattern separation via binwise compactness, binwise occupancy or FR codes were measured for each recording set of every celltype, as in Fig 6B (negative values indicate pattern convergence). Levels were different between celltypes (Kruskal-Wallis one-way ANOVA: p < 0.0001 at all time scales; post-hoc t-tests with Tukey-Kramer correction for multiple comparisons in **Table 4**).

Overall, we demonstrate here that not all hippocampal celltypes exhibit the same computations, but that they all can perform temporal pattern separation to different degrees through different neural codes and over different time scales. This suggests that different types of neurons can serve different, complementary, computations that allow the isolated hippocampal network to perform temporal pattern separation using multiplexed coding strategies.

## Discussion

In summary, we report that the hippocampus can perform temporal pattern separation or convergence at the level of single neurons through different multiplexed neural codes. Pattern separation can be achieved by varying many different spike train features in the ensemble of output spike trains, like the spike times, the firing rate, the number of bursts or the number of spikes in a burst. We found that GCs, the output neurons of the DG, mostly use a “desynchronization” strategy leading to an orthogonalization of their output spike trains, especially at short time scales. When considering a neural code focused on firing rates or burstiness, GC output varies little, leading to pattern convergence or separation depending on the degree of similarity of the inputs. Other hippocampal neurons perform different computations through complementary combinations of various neural codes (mostly rate for FSs, burstiness for HMCs and synchrony at long time scales for CA3 PCs), and their network interactions constrain the ability of the hippocampus to perform temporal pattern separation, notably through inhibition of GC activity. Given that, during behavior, hippocampal neurons have a mixed selectivity to different aspects of the environment and cognitive task contingencies [56-58], the use of multiplexed codes is likely critical to encode rich representations efficiently [59]. Therefore, the multiplexed forms of pattern separation we have characterized here might be essential to optimally disambiguate between similar multidimensional memories.

### The hippocampus performs multiplexed forms of pattern separation

Overall, our study highlights that measuring neuronal computations implies a certain vision of the neural code, and that brain networks or single neurons perform many different computations depending on the assumed coding scheme.

The “neural code” is a pervasive but vague concept that refers to the set of features of neural activity that convey information, for example to represent our experiences and memories. The first difficulty in breaking this code comes from the infinitely large number of potential features and combinations of features that could be relevant in the brain. Since the early work of Adrian [60], the most common view is that neurons transmit information about their stimulus through their firing rate, i.e. the number of output spikes averaged over a certain time window. This rate code is often contrasted to time codes, but this is a false dichotomy [45]: there is an infinite number of possible rate codes, as there is an infinite number of time windows over which the number of spikes could be integrated (are spikes counted by reader neurons over milliseconds, seconds or minutes?). In addition to spike counts, information can also be carried by the timing of single spikes [45, 46, 61], bursts [62, 63] or even more complex spike patterns [61, 64-66]. Even when focusing on time or burstiness codes, many subfeatures could be considered words in the elusive language of the brain: for example, the information could be encoded in the specific spike times [67] or the interspike intervals [64, 68], the relative or exact latency or synchrony with regard to other spike trains [67] or the relative timing of individual spikes with regard to a reference network oscillation [69], the timing or frequency of bursts [62, 63], the spike times within a burst or the spike counts per burst [63, 70].

Pattern separation has long been thought to cover several potential computations depending on the neural code considered [5]. Indeed, past investigations did not always define brain activity patterns the same way, nor did they choose the same similarity metrics and time resolution, thus assuming different forms of pattern separation. When the hypothesis of pattern separation was first formulated, activity patterns were defined as binary vectors representing populations of ON or OFF neurons, without considering a time dimension [4, 6, 44]. As a result, modeling work that followed often excluded the dynamics of neural activity, and used the number of common active neurons between two population patterns as their measure of similarity [43, 54, 71]. In addition to this purely spatial population code, others have defined activity patterns as maps of firing rates averaged over minutes, or as population matrices of those maps [12, 14], measuring pattern separation using R [72] or NDP [73] as their similarity metric. Only two modeling studies have investigated pattern separation by directly considering spike trains at a time scale less than a minute: one considered an input pattern as a static population of ON/OFF neurons, an output pattern as a population vector of firing rates averaged over 500 ms and the similarity between two output patterns was based on the sum of firing rate ratios [71]; the second considered an input pattern as an ensemble of 30 ms spike trains (0 or 1 spike per input channel, i.e. akin to a population of ON/OFF neurons but with the time of an input spike carrying some information due to the possibility of temporal summation) and an output pattern as an ensemble of 200 ms spike trains, using R with a τ_w_ of 20 ms to measure the similarity between spatiotemporal patterns [41].

Past computational studies suggest, together, that the DG could perform pattern separation through different codes, but our investigation is the first to systematically explore and compare pattern separation levels based on diverse coding strategies. It is also the first focusing on time codes. Thanks to our slice-based pattern separation assay, we could control input patterns while recording the output of the DG, thus allowing us to experimentally test pattern separation in the DG for the first time [5, 40]. We considered activity patterns (for both input and output) to be two-second long spike trains, and used multiple definitions of spike train similarity at time scales going from submillisecond to second resolution. We first measured spike count correlations with R and NDP, two common metrics with different sets of assumptions on the neural code (e.g. NDP does not consider common periods of silence as correlated; see **Supplements**), showing that the DG performs decorrelation and orthogonalization of its outputs at subsecond time scales at the level of single GCs (**Fig 3A-B**). Because this phenomenon is stronger at millisecond time scales, and that R and NDP are mostly sensitive to binwise synchrony, we measured spike train similarity with SPIKE, assuming a synchrony code purely based on spike times instead of spike counts [48]. This confirmed that GC outputs are separated through “desynchronization” of their spike times from sweep to sweep (**Fig 3D**). Moreover, we designed a new simple similarity metric, SF (which complements NDP in the description of the similarity between two spike count vectors), as well as two complementary measures of spike train “burstiness” (compactness and occupancy). These revealed that, in addition to desynchronization strategies, GCs can exhibit pattern separation through firing rate or burstiness codes, and that depending on the time scale, the neural code considered and the input statistical structure, different levels of pattern separation or convergence are performed (**Fig 6**).

In conclusion, our work demonstrates that single neuron pattern separation can be achieved in isolated hippocampal tissue through multiplexed coding, which is a way of simultaneously representing several features of a stimulus or, in our case, simultaneously performing different computations, through different temporal scales and spike train features [59, 63, 74]. Our work illustrates the importance of considering multiple neural codes when studying neuronal computations for three reasons: 1) The fact that conceptually different metrics lead to converging results bolsters our conclusion that temporal pattern separation occurs within the DG and CA3 at the level of single neurons; 2) a single neuron can exhibit either pattern convergence or separation depending on the coding scheme; 3) a given neural code can be more relevant for pattern separation/convergence in certain celltypes than others.

Our finding that DG can perform pattern convergence in some conditions is particularly intriguing. If pattern separation is conceptualized to support mnemonic discrimination, then pattern convergence would support generalization, another elusive mnemonic phenomenon [75]. In the future, experiments correlating mnemonic discrimination and generalization with the levels of pattern separation or convergence through various potential codes will help pinpoint which computations are actually used in the brain to support episodic memory.

### Network mechanisms supporting hippocampal computations

Our work, in showing that spiking was difficult to elicit in GCs and that their responses were sparser than their inputs (**Fig 5**) is in line with past research [20, 23]. It also resonates with the long-standing idea that the DG acts as the gate of the hippocampus, filtering out incoming cortical activity before it reaches CA3 to prevent overexcitation [49]. Figuring out the computations of the DG is equivalent to reverse-engineering the transfer function defining the DG filter. Some have suggested that the DG is a low-pass [22, 76] or a ∼10 Hz band-pass filter [21]. Our results confirm that the output frequency of GCs is constrained in a narrow band (**Fig 5B**) and go one step further by providing the first demonstration that the filtering properties of the DG allow it to perform different computations including pattern separation in response to complex naturalistic inputs (**Fig 6**).

The question is then: how are these filtering properties implemented? The physiological mechanisms underlying DG computations in general are understudied and remain a mystery. In Madar et al. (2018) [40], we provided a proof-of-principle that the short-term dynamics of synaptic transmission at the perforant path-GC synapses could explain the levels of temporal pattern decorrelation in the DG. But GCs also receive feedforward and feedback inhibition and excitation from multiple interneurons that also interact between each other [51, 77, 78]. Indeed, our finding that partial block of GABA_A_-mediated neurotransmission impairs temporal pattern separation in the DG (**Fig 7**) strongly suggests that the computations we investigated are network phenomena. Interactions between multiple celltypes are likely involved, and it is thus critical to characterize the suprathreshold input-output transformation of each celltype to parse out their computational role. In the present work, we shed light on the computations of dentate FSs (also known as basket cells providing fast feedforward and feedback perisomatic inhibition to GCs [79]), HMCs (glutamatergic interneurons providing feedback monosynaptic excitation and disynaptic inhibition to GCs mostly outside of the lamellar plane [77]), and CA3 PCs, the output neurons of the hippocampal subnetwork directly downstream of the DG in the trisynaptic circuit, but which can also provide indirect feedback to the DG via collaterals targeting hilar neurons [77]. We discovered that, in contrast to GCs (**Fig 8-9**, and see [40]): 1) FSs generally exhibit low levels of pattern separation through binwise desynchronization (R, NDP) or scaling (SF, likely due to high firing rates), but high levels of separation through simple variation of local firing rates from sweep to sweep, especially at the 100 ms time scale and higher; 2) HMCs exhibit intermediate to low levels of binwise desynchronization up to the 500 ms time scale. However, contrarily to GCs and FSs, they are able to achieve high levels of pattern separation through scaling, thanks to a variable bursting response and low overall firing rates; 3) CA3 PCs exhibit high levels of pattern separation through all tested neural codes, but significantly more than their GC controls through binwise desynchronization only, especially at long time scales, suggesting that CA3 might complement and amplify the separation inherited from the DG, possibly to make pattern completion more efficient [40]. In the future, computational modelling combining those results with knowledge of the synaptic dynamics between these populations of neurons will help dissect the role played by each element of the network in hippocampal computations.

Our study especially calls for more work on the computational role of inhibitory synaptic dynamics as it reveals the first experimental evidence that fast GABAergic transmission favors pattern separation through multiple neural codes (binwise desynchronization and scaling) and impairs it through another code (occupancy variations at short time scales) (**Fig 7**). Past research has long suspected the importance of inhibition to hippocampal computations. Computational models of the DG implementing perisomatic feedforward inhibition (provided by FSs) in the form of a “k-winners take all” rule controlling the number of GCs activated by a given input pattern have stressed the importance of such inhibition on pattern separation through a population code [4, 54, 55]. The seminal model of Myers and Scharfman (2009) also suggested that certain hilar interneurons, providing feedforward and feedback inhibition on the distal dendrites of GCs, could impair population pattern separation through their influence on other GABAergic interneurons (see also Danielson et al. 2017). Experimentally, a recent behavioral study on mice showed that normal levels of tonic inhibition in GCs are critical for mnemonic discrimination [80]. Our results now demonstrate that inhibition is also critical for computations that could support this episodic memory function. Future experiments will need to clarify the exact role of different interneuron populations as well as different types of inhibition (e.g. feedforward vs feedback or somatic vs dendritic). Such work will especially be critical to further understand CA3 computations and how they differ from those of DG, because our findings on CA3 are based on recordings with partially reduced inhibition. On one hand, the enhancement of pattern separation we observed in CA3 could be due to abnormal pharmacological conditions, on the other hand, the necessity to lower inhibition to get CA3 PCs to spike could instead indicate that CA3 must be disinhibited during episodic memory formation.

## Methods

Herein, we revisited experiments performed in Madar et al. (2018) [40] and expanded on them by using additional types of input patterns, new pharmacological conditions, and by analyzing the previous and new datasets with multiple similarity metrics assuming different neural codes.

### Experiments

All experiments were performed in accordance with the National Institute of Health guidelines outlined in the National Research Council Guide for the Care and Use of Laboratory Animal (2011) and regularly monitored and approved by the University of Wisconsin Institutional Animal Care and Use Committee.

Briefly, we performed experiments on C57Bl6 male mice (Harlan/Envigo) following the general paradigm described in Madar et al. (2018) [40]. Horizontal acute slices of a mouse brain hemisphere containing the hippocampus were prepared in cutting solution (CS; two version were used. CS#1/CS#2, in mM: 83/80 NaCl, 26/24 NaHCO_3_, 2.5/2.5 KCl, 1/1.25 NaH2PO_4_, 0.5/0.5 CaCl_2_, 3.3/4 MgCl2, 22/25 D-Glucose, 72/75 Sucrose, 0/1 Na-L-Ascorbate, 0/3 Na-Pyruvate, bubbled with 95% O_2_ and 5% CO_2_) and recorded at 33-35°C in regular artificial cerebrospinal fluid (aCSF, in mM: 125 NaCl, 25 NaHCO3, 2.5 KCl, 1.25 NaH2PO4, 2 CaCl2, 1 MgCl2, and 25 D-Glucose). Two second input trains of varying similarity were delivered to neural afferents of DG or CA3 neurons via bipolar electrical stimulation through a glass theta pipette, while recording the membrane potential of a single responsive patched neuron in whole-cell current-clamp mode. For the intracellular solution (IS), we used two different recipes that yielded similar electrophysiological behaviors and results (IS#1/IS#2 in mM: 140/135 K-gluconate, 0/5 KCl, 10/0.1 EGTA, 10/10 HEPES, 20/20 Na-phosphocreatine, 2/2 Mg_2_ATP, 0.3/0.3 NaGTP, 0/0.25 CaCl2, adjusted to pH 7.3 and 310 mOsm with KOH and H_2_O). The location of the stimulation (>100um away from dendrites of the recorded neuron) and its intensity (0.1-1 mA) was determined to yield ∼50% probability of spiking after an input pulse (range: ∼20-80%). A stimulation protocol consisted of five or ten trains (i.e. a given input set), interleaved (with 5 s of pause between each train) and repeated ten or five times, respectively, such that fifty output traces were recorded (**Fig 1C**). The amount of pattern separation or convergence effectuated for a given recording was assessed by pairwise similarity analysis (defined below), displayed as what we refer to as a “pattern separation graph”: the pairwise input similarity across all pairs of input trains (excluding self-comparisons) was compared to the average similarity of all pairs of output spike trains coming from the corresponding pairs of input trains (excluding comparisons between output spike trains coming from the same input train) (**Fig 1D-E**).

**Figs 1-3** (GCs) and **Figs 8-9** (all celltypes) are based on recordings from GCs, fast-spiking interneurons (FSs), hilar mossy cells (HMCs) and CA3 pyramidal cells (CA3 PCs) reanalyzed from Madar et al. (2018) [40]. They were performed on slices (CS#1 for GCs, CS#2 for other celltypes; IS#1 for all) from young mice (p15-25), using 11 input sets consisting of five Poisson spike trains with a mean rate of 10 Hz (GCs, FSs, HMCs) or 30 Hz (CA3 PCs and their GC controls). GCs (10 Hz), FSs and HMCs were recorded in regular aCSF. Because it was very difficult to promote firing of CA3 PCs in aCSF, CA3 PCs and their GCs control were recorded in aCSF with 100 nM gabazine (gzn, SR-95531) to decrease inhibitory postsynaptic currents (IPSC) amplitude by ∼30% and allow CA3 PCs to occasionally escape inhibition and fire action potentials. Note that the GCs recorded in gabazine as controls for the CA3 PC experiments could not be directly compared to GC recordings in regular aCSF because input sets with different mean rates were used.

To determine the role of inhibition on pattern separation in GCs, we performed additional recordings of single GCs in slices from young mice (p22-32, 6 animals) (**Fig 7**). After cutting (CS#2), slices were moved to a storing chamber containing 100% CS, at 37°C for 30 minutes after dissection. As in all other experiments, the storing chamber was then placed at room temperature and slices left there for at least 30 min. Slices were then transferred to the recording chamber and perfused (5 ml/min) with oxygenated regular aCSF at 33-35°C. Patch-clamp recordings of GCs were performed using IS#1. An experiment consisted of recording the same GC under the same input set of five 10 Hz Poisson trains (average R_input_ = 0.76 at 10 ms, shown in **Fig 1B**) before and after addition of 100 nM gabazine to the flow of regular aCSF. We waited at least five minutes after the solution switch, to allow for equilibrium to be reached. The holding potential (∼-70mV), the location of the stimulation electrode, the stimulus current intensity and all other parameters were unchanged between the two recordings (e.g. under gabazine, the spiking probability was not retargeted to ∼50%, unlike the gabazine experiments using 30 Hz inputs mentioned above). The intrinsic properties of recorded GCs (n = 7) were (mean ± SEM): 1) Before gzn: resting membrane potential V_rest_ = −76.5 ± 2.5 mV, input resistance R_i_ = 194 ± 30 MΩ and membrane capacitance C_m_ = 20 ± 1.4 pF; 2) After gzn: V_rest_ = −74.4 ± 2 mV, R_i_ = 184 ± 29 MΩ and C_m_ = 17 ± 1.6 pF. R_i_ was not significantly affected and V_rest_ and C_m_ were only slightly decreased after treatment (Wilcoxon signed rank test: V_rest_ p = 0.09; R_i_ p = 0.44; C_m_ p = 0.03). The slight drift of C_m_ was likely due to the small increase in access resistance occurring over time (before: 6 ± 0.8 MΩ, after: 7.3 ± 1.4 MΩ; p = 0.06).

To explore the impact of a variety of input pattern statistical structures on single neuron computations, we designed two new input sets comprised of 10 different two seconds spike trains (**Fig 4-6**). Trains in “input set A” follow a Poisson (P) distribution and have an average Pearson’s correlation of 75% at the 10 ms time scale, but each train has a different mean firing rate (PΔFR, **Fig 4A**). Trains in “input set B” do not follow a Poisson distribution: they all have 21 spikes, making them all 10.5 Hz overall, but are more or less bursty (B) (i.e. for each spike train the same number of spikes was distributed in a different number of time bins. Occupied time bins were selected randomly – following a uniform distribution – among all the possible time bins) (B 10.5 Hz, **Fig 4B**). For these experiments, we used slices from two adult mice (p145 and p153) injected with saline at p46 as controls for a separate set of experiments (not described here). Slices were prepared as described above (CS#2) and patch-clamp recordings were performed in regular aCSF using IS#2. The intrinsic properties of recorded GCs were: 1) Input set A (n = 5): V_rest_ = −80.4 ± 2.9 mV, R_i_ = 155 ± 19 MΩ and C_m_ = 23 ± 2 pF; 2) Input set B (n = 3): V_rest_ = −78.0 ± 5.5 mV, R_i_ = 178 ± 14 MΩ and C_m_ = 24 ± 3 pF.

All chemicals and drugs were purchased from Sigma-Aldrich (USA).

### Input spike trains

As explained above, three types of input sets were used, defined by the statistical structure (P for Poisson, B for Bursty) and the firing rate of their spike trains (constant or varying across trains): 1) P10Hz (**Fig 1-3, 7-9)**, 2) PΔFR (input set A in **Fig 4-6**), 3) B10.5Hz (input set B in **Fig 4-6**).

Input sets with Poisson spike trains (P10Hz and PΔFR) were generated using two different algorithms allowing to specify the number of spike trains in an input set (5 or 10, respectively), the firing rate of each spike train, and the average Pearson’s correlation coefficient across all spike trains (computed with τ_w_ = 10ms). First, through iterative modifications of a random Poisson spike train, we generated five different P10Hz input sets of 5 trains with average correlation R_input_ = 0.88, 0.84, 0.74, 0.65, 0.56. Six other P10Hz input sets of 5 trains (R_input_ = 1.00, 0.95, 0.76, 0.48, 0.26, 0.11) and one PΔFR set of 10 trains (R_input_ = 0.75) were generated using the Matlab toolbox provided by Macke and colleagues [81], which uses a more efficient and mathematically principled algorithm that randomly samples from a multivariate Poisson distribution with preset rate (FR) and covariance matrix. For both algorithms, although pairwise correlations were constrained around the specified mean R_input_ (e.g. at τ_w_ = 10 ms, on average across all P10Hz input sets, the relative standard error of pairwise R_input_ values is 4% of the mean R_input_), some variability remained (see **Fig 1E right**) which allowed to test a wide array of pairwise input correlations (**Fig 1F**). Because this variability may be larger when measuring spike train similarity with different metrics than R, or at τ_w_ values that were not used to design input sets, we reported pairwise similarity values (as opposed to the average similarity across all trains) throughout the article.

To design an input set with 10 spike trains spanning a range of burstiness (B10.5Hz), we simply constrained the number of spikes to be equal in all trains and specified the number of bins with at least one spike for each train (bursty trains have a lower number of occupied bins). Spikes were assigned to randomly selected bins and to random times within those bins.

### Similarity metrics

We assessed the similarity between spike trains in four ways, spanning a range of different assumptions on the neural code: 1) with the Pearson’s correlation coefficient (R), 2) with the normalized dot product (NDP), 3) with the scaling factor (SF) and 4) with a distance metric called SPIKE specifically designed to assess the dissimilarity between two spike trains [48]. In addition, we used measures of differences between spike trains that assume different neural codes, related to the one assumed by SF but easier to interpret in terms of basic spike train features (see **Firing rate and burstiness codes** and **Dispersion metrics** sections below).

For R, NDP and SF, two spike trains X and Y of the same duration were divided into N time bins of duration τ_w_, with X_i_ and Y_i_ the respective numbers of spikes in bin i for each spike train. In contrast, the SPIKE metric is binless and based on the spike times of each spike train. For all similarity measures, empty spike trains (i.e. sweeps during which no spikes were evoked) were excluded.

R (Pearson’s correlation coefficient) can take values between 1 (perfectly correlated, i.e. identical) and −1 (anticorrelated) with 0 indicating X and Y are uncorrelated. It was computed with the following equation, where *cov* is the covariance (scalar), σ is the standard deviation and 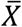 and 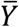 are the means of X and Y:

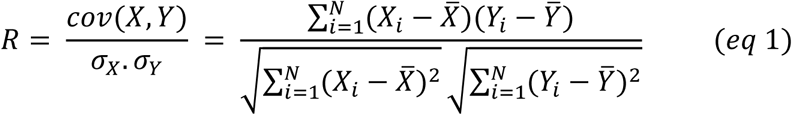

NDP (Normalized Dot Product) is the cosine of the angle θ between the two vectors: it is 0 when they are perfectly orthogonal, 1 when they are collinear (see **Fig 2A, C**). NDP was computed as the dot product of X and Y divided by the product of their norms, with the following equation (where X_i_ and Y_i_ are the coordinates of X and Y):

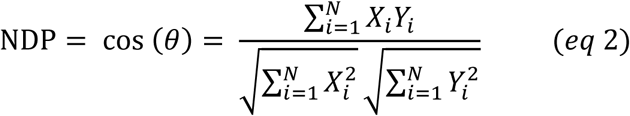

SF (Scaling Factor) quantifies the difference of length between the two vectors X and Y. We have defined it as the ratio between the norms of each vector, the smaller norm always divided by the bigger one to have SF values ranging from 0 to 1. SF = 1 means X and Y are identical in terms of binwise spike number. The closer to 0 SF is, the more dissimilar are X and Y (see **Fig 2A, C**). SF was computed with the following equation (where 0 < ||X|| ≤ ||Y||):

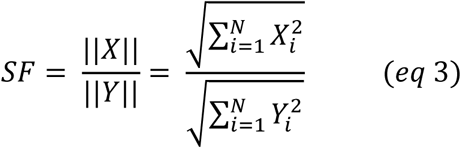

NDP and SF focus on complementary features of a system of two vectors (angle vs norms), and thus, together they are sufficient to fully describe the similarity between two vectors in a Euclidean space.

The binless SPIKE similarity metric was computed from lists of spike times using the Matlab toolbox provided at **www.fi.isc.cnr.it/users/thomas.kreuz/sourcecode.html**. The provided software computes the SPIKE-distance between two spike trains (called D(t) in our study and S(t) in eqn. 19 of the original paper) [48] and we derived the SPIKE similarity as: 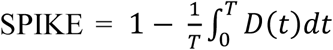, where T is the duration of the spike train. Because D(t) ranges from 0 to 1, SPIKE is thus also between 0 and 1, like NDP and SF. When SPIKE is equal to 1, spike trains have exactly the same spike times, i.e. they are synchronous (n.b. in our experiments, spike trains were not simultaneously recorded, but we use “synchronous” in the sense of spike trains aligned to the start of each sweep). Note that SPIKE has a large dynamic range (i.e. sensitivity over large differences of spiketimes), and, as a result, realistic spike trains like in our input sets rarely have a SPIKE similarity lower than 0.45 [48, 82] (see **Fig 3D**).

The ranges of possible values for the metrics we used are listed below, where closer to 1 always means more similar (although what similar means is different for each metric):

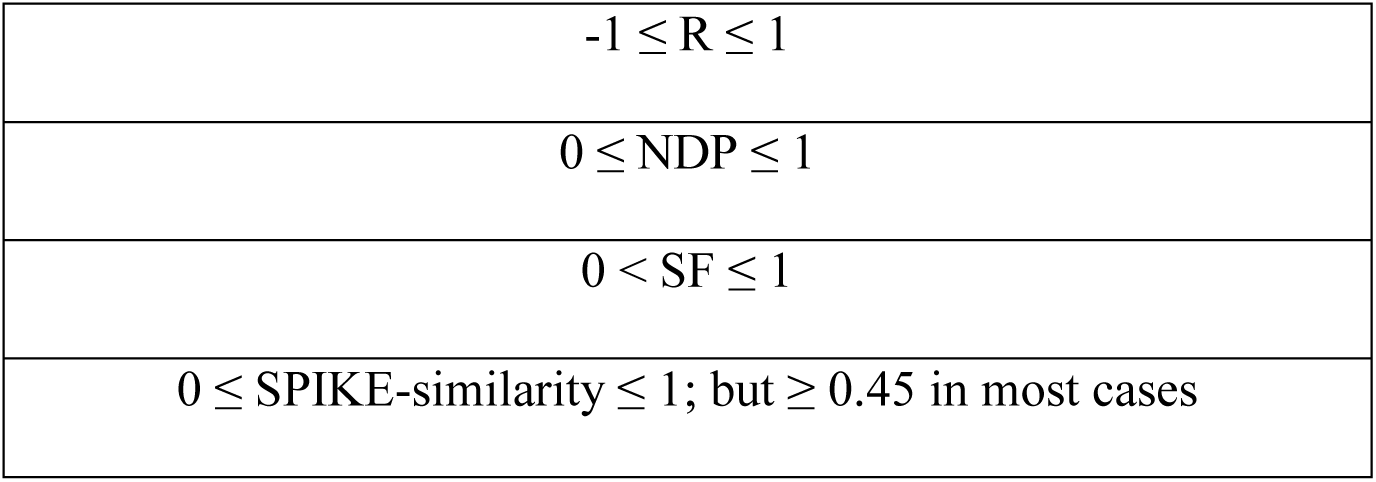

Each metric assumes a specific neural code. For instance, R, NDP and SF all consider that the basic informative feature in spike trains is the number of spikes in a time bin (i.e. a spike count code), but R and NDP suppose that spike trains are similar when the spike counts increase or decrease similarly in the same time bins (with empty bins not counting for NDP), thus assuming a binwise synchrony code, whereas SF assumes a code where it is the similarity in overall burstiness or firing rate that is relevant (see **Fig 4A-B**), not the binwise synchrony (see **Fig 2A-C**). SPIKE, on the other hand, considers a spike time synchrony code. Details on the neural codes assumed by each metric, how they differ from each other and how they are impacted by global increases in firing rate can be found in the **Supplements** (part 1). The main assumptions and properties of each similarity metric on the neural code are summarized in **Table 1**. Note that we do not consider one metric to be better than the other, or better than alternative metrics not used in this study. Each metric can be used as a tool to analyze different aspects of the neural code, with a set of pros, cons and assumptions that complement other methods.

### Firing rate and burstiness codes

R, NDP, SF and SPIKE are considering different aspects of spike trains and thus assume different neural codes. However, it can be difficult to interpret those similarity measures in terms of common properties of spike trains like the firing rate or the burstiness. To get a better sense of the neural code assumed by each similarity metric, and to directly assess whether pattern separation was performed through easily interpretable strategies like variation of the firing rate or variation of the burstiness, we measured such properties in each spike train.

The firing rate is simply the number of spikes divided by the duration of a sweep (2 seconds). Measuring burstiness is less straightforward as there is currently no consensus on the definition of a burst of action potentials [83-85]. We used three measures, detailed below.

Spike trains are often considered bursty when they diverge from a Poisson process [28, 85]. To determine the tendency of a neuron to fire bursts, we computed the burstiness of a given output set as the Kullback-Leibler divergence (D_KL_) of its interspike interval (ISI) frequency distribution M (normalized to the median ISI) from the normalized ISI frequency distribution P of a Poisson process. In practice, output sets from a 10 Hz input set were compared to the ISI distribution from all combined 10 Hz input sets, whereas input sets from 30 Hz input set were compared to the combined distribution of all 30 Hz input sets.

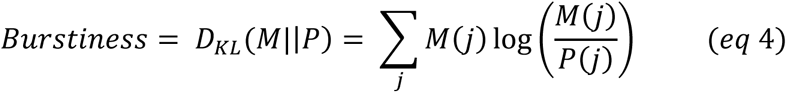

where M(j) or P(j) is the value of M or P in the jth bin of the distribution. Normalized ISI frequency distributions were discretized in 250 bins going from 0 to 50, and D_KL_ was computed using the KLDIV matlab function (v 1.0.0.0), written by David Fass, available at: https://www.mathworks.com/matlabcentral/fileexchange/13089-kldiv.

See **Supplements - 2** for details on why D_KL_ is a suitable measure of cell burstiness.

To assess burstiness in single spike trains, the D_KL_ measure being not well-suited for this (see **Supplements - 2**), we designed two simple nonparametric measures that we called *compactness* and *occupancy*. Instead of explicitly defining “burstiness”, these measures are based on the binning of spike trains, offering the advantage of being directly comparable to the neural codes assumed by binned similarity metrics like R, NDP and SF (**Fig S2**). A spike train is thus seen as a vector of spike counts, and a time bin (or vector coordinate) is considered a window in which spikes can be clustered. Thus, instead of counting the number of detected bursts and the number of spikes in a burst, our two measures provide information on the number of occupied bins (Compactness) and the number of spikes in occupied bins (Occupancy). These measures of burstiness should be viewed as complementary to previous definitions, not as replacements.

The compactness of a spike train was computed as follows:

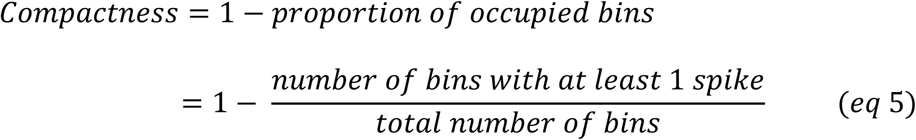

Compactness thus runs between 0 (all bins are occupied by at least one spike) and 1 (no spikes: this case was excluded so the true maximum, corresponding to any spike train with all its spikes clustered in just one bin, was 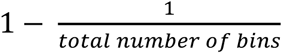). Values closer to 1 mean very few bins are occupied: either many spikes are clustered in a small number of bins or the measured spike train has very few spikes.

The occupancy of a spike train is the average number of spikes per occupied bin:

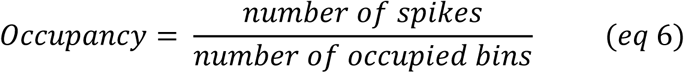

Because we excluded empty spike trains, the occupancy can theoretically go from 1 to infinity. Note that the lowest value (i.e. 1 spike per occupied bin) means that all spikes are distributed in different bins (or that there is just 1 spike).

Compactness and occupancy are both dependent on the firing rate of a given spike train, but not in the same way (see **Supplements - 2** for details). They are complementary measures that, together, allow determining whether two spike trains with different firing rates are also different in terms of burstiness (**Table S1**). When spike trains have the same firing rate (e.g. input set B 10.5 Hz in **Fig 4B**, which were all spike trains with 21 spikes each but were constructed by specifying different numbers of occupied bins), compactness and occupancy are redundant (**Fig S2C-D**) and, for both measures, higher values mean higher burstiness.

### Dispersion metrics

To assess the amount of pattern separation performed by variation of a given spike train feature like the firing rate, compactness or occupancy, we measured the difference in this feature between two spike trains. In the same spirit as for similarity metrics, we could then compute the difference between all pairs of trains in an input set (excluding self-comparisons) and compare to the difference of all corresponding pairs of output spike trains, (excluding comparisons between output spike trains associated with the same input spike train). This would yield matrices of pairwise differences organized in the same fashion as the similarity matrix displayed in **Fig 1E**, which could then be converted to pattern separation graphs showing the average pairwise output absolute difference as a function of the pairwise input difference (**Fig 6A**). Note that the mean absolute difference (MAD) between all spike trains of a given set (either input or output) corresponds to the Gini mean difference [86], a common measure of dispersion of a sample akin to the standard deviation (**Fig 5A**). The difference in MAD between an input set and its output set indicates whether pattern separation or convergence was achieved through variation of the measured spike train feature (**Fig 6B**).

### Software and statistics

Data analysis was performed using MATLAB (Mathworks, Natick, MA, USA). Sample sizes were chosen based on the literature and estimations of the variance and effect size from preliminary data. The one-sample Kolmogorov-Smirnov test was used to verify the normality of data distributions. Parametric or non-parametric statistical tests were appropriately used to assess significance (p-value < 0.05). Assumptions on equal variances between groups were avoided when necessary. All tests on means or medians were two-tailed. To determine whether distributions of similarity values [S_input_, S_output_] (S standing for any similarity metric) were significantly different at a given time scale (**Fig 7, 8**), we performed an analysis of the covariance (ANCOVA) using separate lines regression models, with S_input_ as a continuous predictor and Treatment (**Fig 7**) or Celltype (**Fig 8**) as a categorical predictor with 2 or 5 levels respectively (the *aoctool* function in Matlab and a custom-written code yielded the same results). The 95% confidence interval around the slope and intercept of a given linear model was determined from the *regress* function (**Fig 8**).

### Data and code availability

Data and code used in this manuscript are freely available:

Data: https://www.ebi.ac.uk/biostudies/studies/S-BSST219

Code: https://github.com/antoinemadar/PatSepSpikeTrains

## Supplements

This document provides supplementary Figures, tables and a discussion on how to interpret the similarity and burstiness metrics used in the article “Temporal pattern separation in hippocampal neurons through multiplexed neural codes”.

## 1 How to interpret similarity metrics

Measuring the similarity between two spike trains is a difficult problem [1, 2], and many methods have been designed to this effect, each with a set of pros, cons and assumptions [2-6]. We could not be exhaustive but, in order to span a large range of potential neural codes, we used four different similarity metrics: R (Pearson’s correlation coefficient), NDP (Normalized Dot product), SF (Scaling Factor) and SPIKE (1 - SPIKE distance). (Note: ‘metrics’ is used with a loose sense throughout the paper). Their pros, cons and assumptions are detailed in this section and summarized in **Table 1** of the main text.

### Binned vs binless metrics

We used two different kinds of similarity metrics: 1) binned metrics (R, NDP, SF) consider a spike train as a list X of spike counts X_i_ (with X_i_ the number of spikes in time bin i of a given duration τ_w_) and thus assume a neural code where spike counts are the basic informative feature; 2) a binless metric SPIKE that considers a spike train as a list of spike times and thus assumes a neural code where the time relative to the start of the given spike train is the basic informative feature.

Although R has already been used to measure the similarity between spike trains [2, 7], none of the binned metrics we used were designed with spike trains in mind and thus have properties that might not be well suited for the study of spike trains. For instance, R, NDP and SF have the drawback of considering all X_i_ (i.e. binned spike counts) as independent observations, which may not be a realistic assumption. Moreover, they assume spike trains are related linearly, or that they belong to a Euclidean space. Binless metrics like SPIKE avoid these limitations [8].

Another limitation of binned metrics as implemented here (and this is also true for compactness and occupancy) is that they are subject to edge effects: small shifts of the bins can include or exclude spikes, which would lead to differences in the similarity measure. However, the fact that marginally different binsizes (e.g. 5 ms vs 10 ms) lead to similar results (in other words, similarity measures as a function of τ_w_ does not show spurious values: see Fig 3A-C, 6A, 7D, 8) suggests that edge effects do not impair our conclusions.

### Differences between binned metrics

#### R vs NDP and SF: epistemological differences

The Pearson’s correlation coefficient R was designed by early statisticians as a tool to quantify the strength and direction of a linear regression [9]. R thus considers two spike trains X and Y as variables, with X_i_ and Y_i_ the observations, and assesses, across all i, whether X_i_ and Y_i_ tend to vary in a similar fashion (**Fig 2B**).

On the other hand, NDP and SF are metrics coming from the fields of Euclidean geometry and linear algebra. In contrast to R, they explicitly consider spike trains as vectors, with each time bin i corresponding to a coordinate, or dimension, in an N-dimensional space (N being the total number of bins). NDP measures the angle between two vectors, and SF is based on vector norms (**Fig 2C**).

#### R, NDP and SF are not equivalent

Although they have been designed for different purposes, both R [7] and NDP [10, 11] have been used to measure the similarity between neuronal activity patterns, including spike trains. In this context, NDP has sometimes loosely been called a measure of correlation [10, 12]. What difference is there between R and NDP then, especially in terms of the assumed neural code? A comparison of equation 1 and 2 (**Methods**) reveals that R is actually an NDP, but of baseline-subtracted vectors. In other words, R does not consider spike trains as raw spike count vectors but as vectors of deviation from their respective mean spike count. This ensures that R is an optimal estimator of correlation [9] but it has the inconvenience of not conserving the angular relationship between X and Y, making NDP necessary if one wants to evaluate pattern separation as an orthogonalization process. Moreover, using baseline-subtracted vectors also means that R considers common silent periods (i.e. X_i_ = Y_i_ = 0) as correlated, which is an assumption on the neural code that has been considered unrealistic by some [2]. In contrast, NDP excludes those empty bins and thus assumes a neural code where common periods of inactivity are irrelevant.

Comparing equations 1, 2 and 3 (**Methods**), the relationships between R, NDP and SF appear not trivial and not linear. To further evaluate the relationship between R, NDP and SF, we compared the values of each similarity metrics for all pairs of a set of simulated spike trains. This set contained the 3^6^ = 729 possible spike count vectors made of six bins, with each bin containing 0, 1 or 2 spikes (some examples are displayed in **Fig 2A**). This analysis confirmed that the three metrics are not equivalent and provides an intuition on what each metric represents (**Fig S1A**).

Interestingly, even though R, NDP and SF are not linearly related in principle, our measures of R and NDP between experimentally recorded spike trains have a near linear relationship, as opposed to R and SF (**Fig S1B**). This correlation is likely due to our choice of input spike trains as well as to biological constraints (e.g. refractory period) forcing the temporal structure of output spike trains to vary only along certain dimensions. It is thus reflective of the actual computations performed in the DG and shows that although R and NDP carry different sets of assumptions about the neural code, spike trains recorded in GCs vary mostly along properties that both R and NDP are sensitive to.

#### R and NDP vs SF: sensitivity to individual firing rate differences and dependency on pairwise firing rate levels

To interpret R, NDP and SF and their different assumed neural codes, it is important to understand their relationship with the firing rate. Studying the impact of firing rate on similarity metrics can however mean different things.

First, a similarity metric can be sensitive to differences of firing rate between two spike trains X and Y. For instance, if X = kY (with k an integer > 1), that means that X and Y have spikes in exactly the same bins (i.e. X and Y have the same temporal structure) but that Y has a higher firing rate. In such a case, both R and NDP would consider X and Y as perfectly similar (see example #2 in **Fig 2A-C**). In contrast, SF is designed to extract 1/k and thus strongly picks up on differences of firing rate. On the other hand, if X and Y have the same firing rate but that their spikes are in different bins (different temporal structure), R and NDP will consider those spike trains completely dissimilar, and SF will see them as perfectly similar (see example #1 in **Fig 2A-C**). Other cases, where X and Y have spikes in the same time bins but only some bins have a different number of spikes (e.g. example #3 in **Fig 2A-C**) reveal that R and NDP are not totally insensitive to differences of firing rate when those are not proportional, but overall these two metrics are mostly sensitive to binwise synchrony, unlike SF (**Fig S2**).

Second, similarity metrics can be influenced by the overall firing rate level of a pair of spike train (for instance measured as the geometric mean of the firing rate of X and Y). Whether an ideal similarity metric should be independent of the firing rate in such a way is debatable, and finding one is not trivial [2, 13], yet it is generally seen as an undesirable property [2] because it makes the comparison of similarity values between two pairs of spike trains more challenging to interpret [1]. Indeed, for spike trains X and Y with random spike times, the probability of spikes occurring close in time will intuitively be higher if X and Y have higher rates [14], even though their temporal structure would be conserved.

Because R is often used to measure spike train correlations [1], its relationship to pairwise firing rate levels has already been studied. It was for instance demonstrated that, when X and Y are two Poisson spike trains with the same rate λ, the normalization factor and the use of variables centered on their mean (which is, in the case of spike trains, a linear function of the firing rate) makes R mathematically independent of λ [14]. However, in the general case, R has still a complex relationship with the firing rates of both spike trains [13].

To get a better sense of this relationship for R, and study the effect of pairwise firing rate levels on NDP and SF as well, we considered two cases: 1) proportional firing rate increases, and additive firing rate increases.

In the first case, we considered two spike trains A and B such that A = kX and B = kY (with k an integer > 1). Some metrics aiming to measure the similarity between spike train in terms of temporal structure, like the correlation index, have been shown to result in higher values for the (A, B) pair compared to the (X, Y) pair [2]. However, in this case, R, NDP and SF values would be the same for both pairs (A, B) and (X, Y): thanks to the denominator in their respective formula, all three are insensitive to proportional increases in firing rates.

In the second case, we used the set of 729 6-bin spike count vectors described above and generated new sets of related vectors with higher firing rates, such that for each spike train X of the original set there was a vector A in the new set with A_i_ = X_i_ + k when X_i_ > 0, and A_i_ = X_i_ otherwise. In other words, the rate was additively (not proportionally) increased while preserving the temporal structure of the initial spike trains and maintaining constant the difference in the binwise number of spikes between spike trains (e.g. for k = 5, if X = [010101] and Y = [020202], then A = [060606] and B = [070707]. For both pairs, the difference is [010101].). Our analysis shows that such an increase in the firing rate can affect all three metrics, but in different ways (**Fig S1C**): NDP and R can increase or decrease, and on average do not change much, whereas SF systematically increases. A larger increase of the firing rate leads to more variability of the effect size for NDP and R whereas the effect monotonically increases for SF.

SF is thus clearly dependent on pairwise firing rate levels, and R and NDP can be affected somewhat as well. Thus, to directly assess the variability of specific spike train features across a set of spike trains (i.e. individual firing rate, compactness and occupancy) without contaminating interpretations with consideration of pairwise firing rate levels (especially in the case of SF), we used additional metrics detailed in **Methods – dispersion metrics**.

#### Temporal resolution of binned metrics

As binned metrics, R, NDP and SF are dependent on the binsize, i.e. a specified temporal resolution τ_w_. In other words, in equations 1 to 3, X_i_ and Y_i_ are functions of τ_w_. Temporal resolution logically influences the value of metrics measuring similarity in terms of the temporal structure of spike trains, such as R [1] or NDP. However it also influences SF: even though SF is sensitive to differences in firing rate, it is designed to be sensitive to differences in binwise firing rate, which makes it also sensitive to differences in burstiness even when the firing rate is constant (e.g. **Fig 4B** and **Fig S2**).

In **Fig 3**, we evaluated the influence of τ_w_ on our results by varying it between 5 ms and 1000 ms. Note that because our spike trains are two-seconds long, using τ_w_ = 2000 ms would mean the spike trains can be seen as variables with only one observation, or as unidimensional vectors (i.e., scalars) that can only vary by their norm (i.e., firing rate). In this case, R is meaningless, NDP is necessarily 1, which indicates collinearity and SF would directly assess the difference in total spike number (i.e. overall firing rate) between the two spike trains.

**Fig S1.**
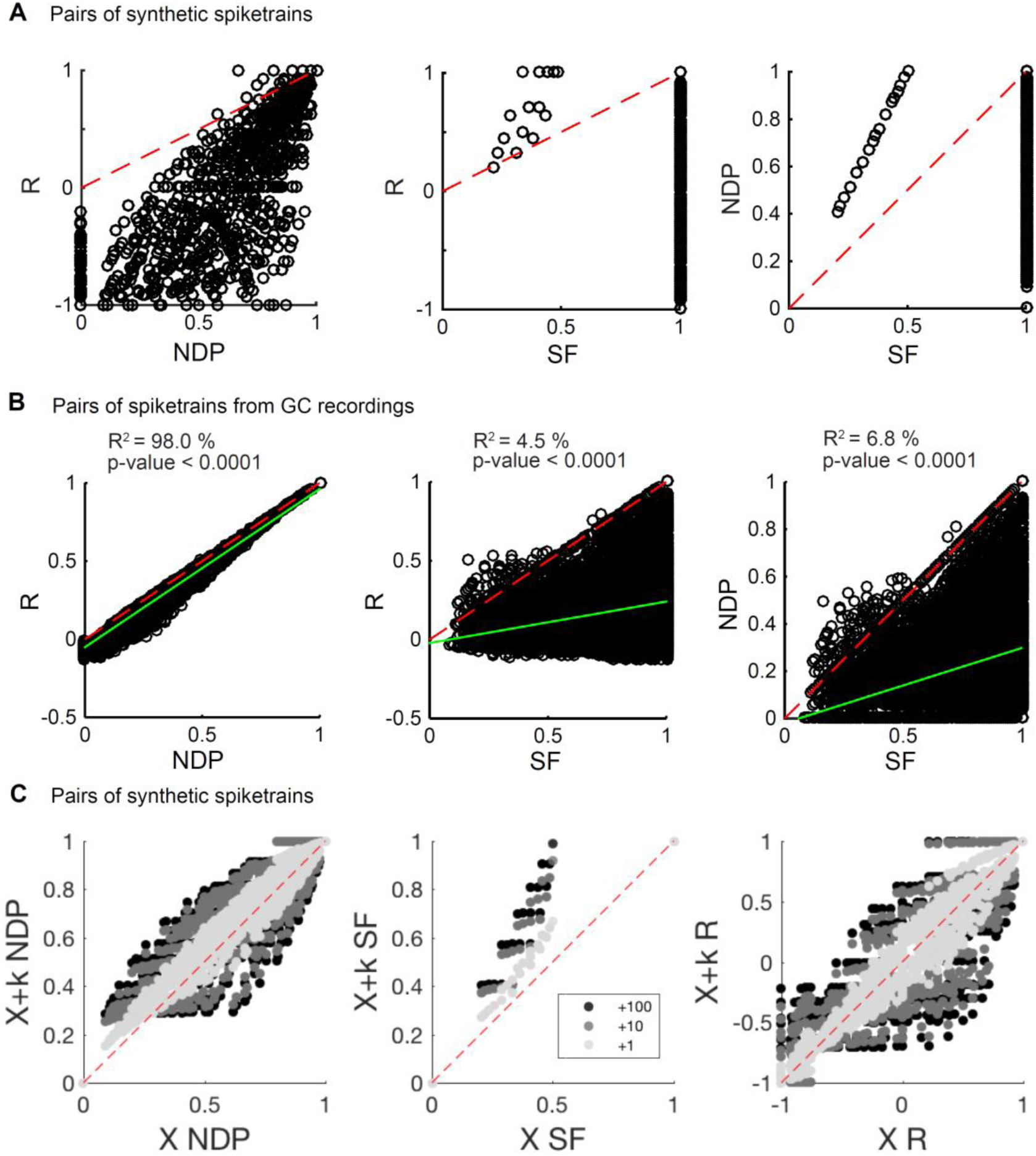
(related to Fig 2). **R, NDP and SF are not equivalent. (A)** Relationships between these 3 similarity metrics (R, NDP and SF) computed between the 3^6^ possible spike count vectors with six bins and that can have 0, 1 or 2 spikes per bin (All pairs combinations = 499,500 data points). **(B)** Relationships between these three similarity metrics for the 124,950 pairs of spike trains from the 102 experimental GC recording sets. Green lines correspond to a linear regression (R^2^ and p-value in each panel). Note that although R and NDP are well correlated in our experimental data (R^2^>0.95), there is not a linear relationship between R and NDP in theory. **(C)** Impact of non-proportional firing rate increase on similarity values for all three metrics. For each 6- bin spike count vector X (same as in A), a vector A was generated such that for each bin i with at least one spike, A_i_ = X_i_ + k, with k = 1, 10 or 100, and A_i_ = X_i_ when X_i_ = 0 (e.g. for k = 1, if X = [010101] and Y = [020202], then A = [020202] and B = [030303]. For both pairs, the difference is [010101]). NDP, SF and R values were computed for all pairs in each set of vectors, and the values of the initial set (x-axis) was compared to the values of the X+k set (y-axis). Data points away from the identity line (red) demonstrate an effect of the firing rate increase.

**Fig S2.**
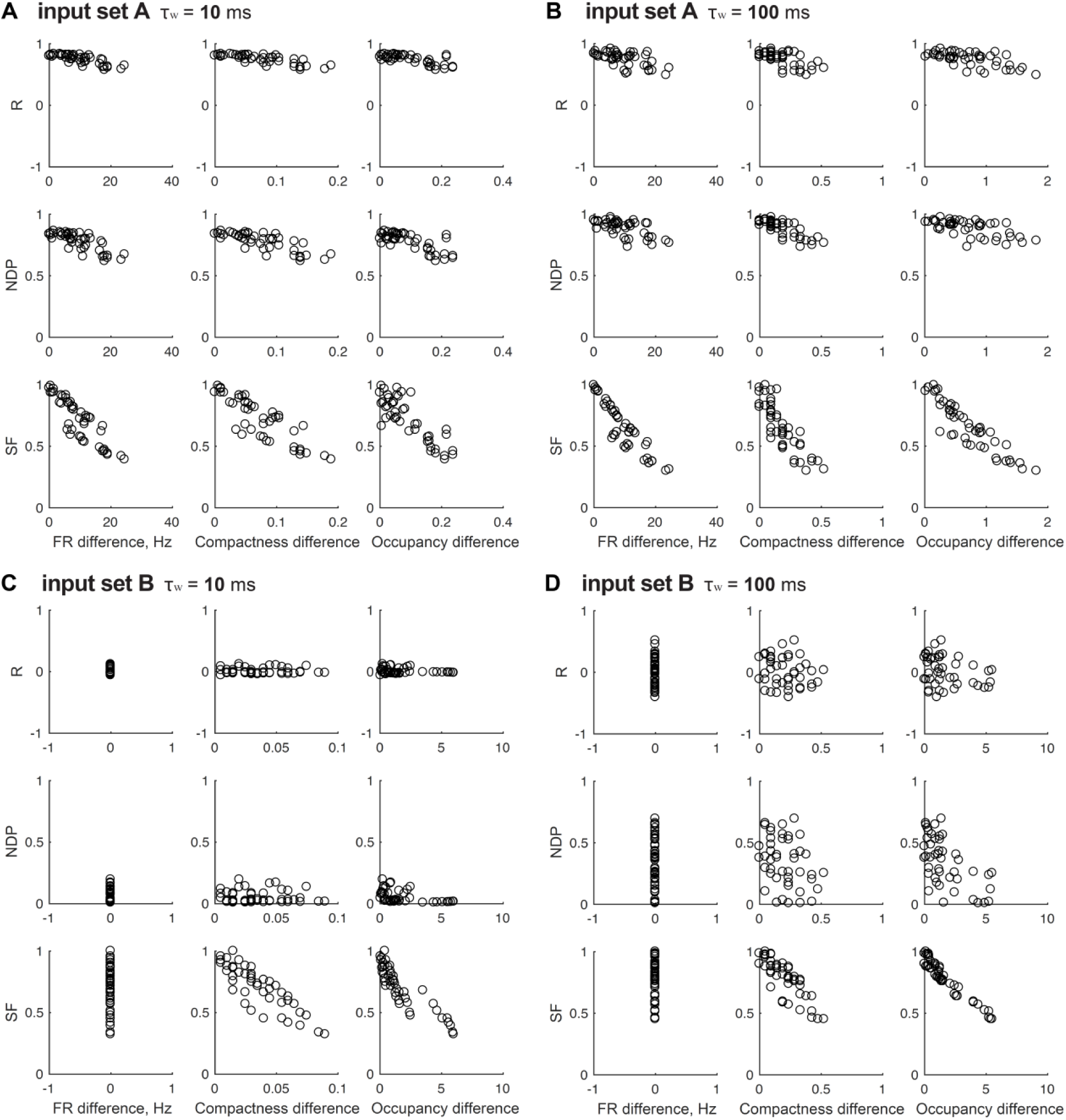
(related to Fig 4)**. Influence of firing rate and burstiness differences on binned similarity metrics.** Data points correspond to single pairs of input trains from input set A (**A-B**) or input set B (**C-D**). The similarity between two spike trains (R, NDP or SF), their difference of compactness (proportion of occupied bins), and their difference in occupancy (average number of spikes per occupied bins) were measured using binning windows of 10 ms (**A, C**) or 100 ms (**B, D**). When FR is constant across spike trains, compactness and occupancy differences are direct measures of burstiness differences, given a certain time scale. As intuited from Fig 2A-C, SF is very sensitive to variations in FR, compactness and occupancy between spike trains, whereas R and NDP are only mildly influenced at best (and mostly at larger time scales), by such variations.

## 2 Burstiness metrics

### Cell burstiness vs spike train burstiness

To assess the tendency of a neuron to fire bursts, we computed the Kullback-Leibler divergence D_KL_(M||P) of the ISI distribution in a set of output spike trains from the ISI distribution of a Poisson process. D_KL_(M||P) is an asymmetric, information theory-based measure of the inefficiency of the assumption that the distribution of ISI M is a distribution P (Poisson, in our case), and thus can be interpreted as the divergence of one probability distribution from another. Note that a train with regularly spaced spikes would diverge from Poissonness without being bursty, but that natural spike trains are rarely regular, and accordingly none of our recordings showed a regular ISI distribution, suggesting that D_KL_(M||P) is a good indirect indicator of burstiness.

The D_KL_ approach above has the advantage of being independent of the firing rate, but has two limitations. First it is parametric, based on the nonconsensual opinion that a Poisson process is the least bursty of processes [15]. Second, because it is based on the comparison of ISI distributions, it requires a large sample of ISIs to be accurate and thus cannot be used to measure the burstiness of a single two second spike train.

In order to assess the burstiness of single spike trains, we designed two binned metrics: compactness and occupancy. We used binned metrics to avoid the difficult problem of burst detection that generally requires some arbitrary threshold to define what is considered a burst [16]. Unlike D_KL_ these metrics are not independent of the firing rate, but together they allow distinguishing between burstiness and firing rate (see below).

### Compactness and occupancy: complementary relationships to the firing rate allow disambiguating between burstiness and lack of sparseness

It is unclear whether the concept of burstiness should be independent or not of the firing rate (for instance, should we consider a spike train with 100 spikes clustered in the same 10 ms bin as bursty as a spike train with just 10 spikes in a single bin?), but, as explained above for similarity metrics, a dependency of burstiness metrics on the firing rate can make interpretations more challenging. Both compactness and occupancy are dependent on the firing rate, but in different ways, which resolves the ambiguity: 1) Compactness does not provide information on the firing rate per se, and is only related to the firing rate in the sense that, in the case of Poisson-like spiking, higher rates lead to a higher chance of occupying a large number of bins. Thus a spike train with just one spike will be considered as compact as a spike train with a large number of spikes all clustered into one bin. In contrast, the occupancy is directly related to the number of spikes and thus to the firing rate of a given spike train (Indeed, when considering a time bin as long as the full duration of a spike train, i.e. two seconds in our case, the occupancy is exactly proportional to the firing rate.). Occupancy would thus distinguish between a sparse spike train with just one spike (Occupancy = 1) and a spike train with 10 spikes clustered in one bin (Occupancy = 10). 2) On the other hand, spike trains with different firing rates can have the same occupancy (e.g. if all spikes are distributed in different bins, Occupancy = 1 regardless of the number of spikes) but would be differentiated by their level of compactness (100 distributed spikes would be considered much less compact than a spike train with just one spike). Therefore, when assessing pairs of spike trains with different firing rates, compactness and occupancy provide complementary information that allows determining whether one spike train is unambiguously burstier or simply has a higher firing rate (see **Table S1**).

**Table S1.** (related to Fig 7) How to interpret compactness and occupancy in terms of burstiness and sparseness when comparing two spike trains? The table shows what it means when *compactness* (C) is increased, stays constant or decreases while *occupancy* (O) increases, stays constant or decreases. We distinguish the cases when the mean firing rate (FR) is constant (FR absolute difference = 0) or different (FR abs diff > 0). The second spike train is either unambiguously burstier (B+) or less bursty (B-), has a greater FR (F+) or is sparser (F-), or a combination of those (i.e. B+F+ or B-F-). In each case, we provide an example (4-dimensional vectors of spike counts) when the conditions can coexist (NA means not applicable). Notice that the resulting table of examples is centrally symmetric.

| C \ O |  | ↗ | — | ↘ |
| --- | --- | --- | --- | --- |
| ↗ | FR abs diff = 0 | **B+**<br>0101 → 0200 | NA | NA |
|  | FR abs diff > 0 | **B+F+**<br>0111 → 0501 | **F-**<br>0202 → 0200 | **F-**<br>0222 → 0101 |
| — | FR abs diff = 0 | NA | 2020 → 0202 | NA |
|  | FR abs diff > 0 | **F+**<br>0101 → 0202 | NA | **F-**<br>0202 → 0101 |
| ↘ | FR abs diff = 0 | NA | NA | **F-**<br>0200 → 0101 |
|  | FR abs diff > 0 | **F+**<br>0101 → 0222 | **F+**<br>0200 → 0202 | **B-F-**<br>0501 → 0111 |

Author contributions
Conceptualization: MVJ, ADM. Data curation: ADM. Formal analysis: ADM, MVJ. Funding acquisition: MVJ, ADM. Investigation/data collection: ADM, LAE. Methodology: MVJ, ADM, LAE. Project administration: MVJ, ADM. Resources: N/A. Software: ADM, MVJ. Supervision: MVJ. Validation: MVJ, LAE, ADM. Visualization: ADM, MVJ. Writing – original draft: ADM. Writing – review & editing: MVJ, ADM, LAE.

## Acknowledgements

We thank Jesse Pfammatter and Kyrie Sellnow for their help in collecting the data.

